# A systematic review and meta-analysis of the effect of experimental pain on rat grimace scale scores

**DOI:** 10.64898/2025.12.22.695961

**Authors:** Xue Ying Zhang, Timothy Pheby, Amber Harris Bozer, Andrew S.C. Rice, Nadia Soliman

**Author notes:** Corresponding author: Nadia Soliman, Pain Research Group, Imperial College, Chelsea and Westminster Hospital campus, 369 Fulham Road, London SW10 9NH, United Kingdom,; URL: https://www.imperial.ac.uk/people/n.soliman16.

## Abstract

The rat grimace scale (RGS) purports to assess behavioural responses related to pain by assessing four facial expressions. To understand how experimental pain affects RGS scores, we conducted a systematic review of studies which reported outcomes using this scale (<u>CRD42022292189</u>). Systematic searches were conducted on 3 databases in January 2022. RGS studies that assessed the effects of disease models relating to pain were included. Risk of bias was assessed by using a modified version of the CAMARADES reporting checklist and SYRCLE Risk of Bias tool. A standardised mean difference effect size was calculated using a random-effects model. Fifty-four studies were included, in which 15 disease model classes and 16 classes of pharmacological interventions were identified. RGS scores were increased in disease models hypothesised to be associated with persistent pain (SMD = −4.44 [95%CI −6.25 to −2.63]) and this response was attenuated by analgesic drugs (SMD = 1.97 [95%CI 1.25 to 2.69]). Post-surgical models (39% of experiments) and opioids (30% of experiments) were the most commonly reported. We could not ascertain the sources of heterogeneity due to insufficient data. Most studies were assessed as having an unclear risk of bias due to poor reporting quality. The relationship between RGS scores and mechanically evoked limb withdrawal was inconsistent. These findings indicate emerging evidence to support the use of RGS in persistent pain research, but additional research is needed to strengthen these findings. Future studies could benefit from using sex-balanced animal populations with diverse genetic backgrounds. To enhance the impact and reproducibility of their findings, studies should prioritise improvements in reporting quality.

## Introduction

Chronic pain is a major global health problem [1], but current pain treatments are limited and have adverse effects. As a result, many patients with chronic pain experience insufficient pain relief. There is a pressing need for more efficacious, safer pain treatments. While promising new analgesics have shown potential in animal studies, they often fail to work effectively in patients [2–5]. There are numerous factors that can contribute to the failure of drugs to advance through the clinical development stages. These factors encompass a range of issues, such as lack of efficacy, safety concerns, a lack of financial incentive to pursue further development, regulatory hurdles that cannot be overcome, problems with manufacturing, legal complications, and development costs or timelines that are impractical. We focus on the halt in the progress of novel analgesic drugs due to their ineffectiveness in clinical trials, despite showing significant efficacy in animal studies. This disparity between results in preclinical and clinical trials regarding a drug’s efficacy has raised concerns about the predictive validity of animal models used to study pain. Researchers are now questioning whether these models accurately represent human pain conditions and if the methods used to measure pain in animals actually reflect pain in humans.

Pain-associated behaviours in animal models are typically assessed using surrogate measures. Among these, stimulus-evoked limb withdrawal is the most common outcome measure [6–9]. However, this has significant drawbacks. Primarily, it lacks real-world applicability as it measures involuntary reflexes rather than the impact of sensory phenotypes associated with disease conditions such as spontaneous pain or sensory loss, which are common in neuropathic pain [3, 10–12]. Additionally, it fails to capture the emotional and physical aspects of pain, which are crucial for understanding the patient experience [10]. Furthermore, animals can learn to anticipate and avoid stimuli, affecting the accuracy of results [13]. Finally, it is difficult to differentiate between pain relief and sedation using the stimulus-evoked limb withdrawal outcome measure, further limiting its reliability. Consequently, the field of animal pain research has been exploring and validating non-evoked outcome measures to overcome these limitations.

Experimental pain has been shown to influence a range of voluntary and learned behaviours in animals [14, 15]. Researchers are increasingly measuring such behaviours to evaluate how persistent and spontaneous pain affects an animal’s physical abilities, emotions, and overall health [16–20]. It is important to recognise that these non-evoked behaviours are not exclusive to pain and therefore require specific interpretation within the context of pain research. To establish a link between these behaviours and pain, studies must demonstrate that changes in these non-evoked behaviours are linked to pain-inducing injuries or diseases and can be alleviated by known analgesics.

Assessing facial expressions in patients with limited verbal communication has shown validity in quantifying pain [21–23]. Based on Darwin’s’ hypothesis of similar facial expressions to emotional states are shared between human and non-human animals [24], grimace scales were developed for mice [25] and rats [26]. Depending on the rodent species, the grimace scale comprises a list of pain-related facial action units that are assessed and scored typically on a scale from 0 to 2. A score of 0 indicates that the human scorer is highly confident that the facial unit is absent. A score of 1 indicates either high confidence in a moderate appearance of the facial action unit or uncertainty regarding its presence or absence. A score of 2 indicates the clear detection of an obvious appearance of the facial action unit with high confidence.

Rodent grimace scores were able to detect pain-related facial expressions in nociceptive assays of moderate duration and was sensitive to known analgesics [26, 27]. However, there are conflicting findings regarding its applicability in persistent pain-related disease models that last hours to days [28, 29]. Moreover, the study of Miller and Leach [30] demonstrated that grimace scores can be influenced by strain, sex, the method of scoring and testing conditions. We conducted a systematic review to only assess the utility and validity of the rat grimace score (RGS). A protocol for a parallel systematic review of mouse grimace score (MGS) has been registered (CRD42021264194).

This systematic review aimed to:

1. assess whether the RGS is affected by experimental conditions relating to pain (i.e., acute nociception and persistent pain-associated disease models) and analgesic drugs
2. explore sources of heterogeneity
3. evaluate the internal validity of studies
4. identify the presence of publication bias
5. assess the correlation between RGS and stimulus-evoked outcome measures

## Materials and Methods

The protocol was registered on PROSPERO (CRD42022292189).

### Search strategy

The systematic search was conducted on 24^th^ January 2022. Three electronic databases were searched: PubMed, EMBASE, and Web of Science.

The search terms were developed around key concepts such as “grimace” and “rats” (Table 1). All studies published until the date of the search were considered without any language restrictions. Duplicates of retrieved studies were removed using EndNote.

**Table 1.** The general search strategy for the systematic review of RGS studies.

| Key Concept | Search Terms |
| --- | --- |
| Grimace Scale | Grimace scale* OR grimace score OR facial grimace |
| AND | AND |
| Rats | Rat OR rats OR rattus OR norvegicus |

### Eligibility criteria for study selection

The inclusion criteria are listed in Table 2.

**Table 2.** Inclusion criteria for the systematic review of RGS studies.

| Component | Description |
| --- | --- |
| **Population** | *In vivo* acute nociception assays and/or rodent models of disease associated with persistent pain.<br>Acute nociception assays assess a healthy rodent’s ability to detect a noxious stimulus, such as the radiant heat induced tail flick assay.<br>Rodent disease models associated with persistent pain are defined as any disease models that are induced chemically, surgically or genetically, developed over a period of hours, weeks or months, leading to pathological conditions associated with persistent pain. A minimum experiment length of an hour. |
| **Intervention** | Any clinically approved or novel drug analgesics used to interfere with nociception |
| **Comparison** | A cohort of control animals:<br>- For acute nociceptive assay and persistent pain associated disease modelling experiments, a control population was defined as<br>i) External control<br>a) A separate cohort of animals that is given a non-noxious stimulus to study acute nociception.<br>b) A separate cohort of sham (prioritised) or naïve animals.<br>c) For studies that used transgenic animals to study persistent pain, a wild-type control was required.<br>ii) Internal control<br>a) A baseline grimace score. It was expected that a proportion of studies may report using baseline grimace scores as the control.<br>- For studies which investigated the effect of analgesic drug interventions on RGS scores, a vehicle control was required. |
| **Outcome** | Studies which assessed rat grimaces as a measure of a pain-related outcome. RGS score data obtained by using the RGS developed by Sotocinal et al. [19]. Studies that used a modified RGS (*i.e.*, *altered facial action units*) were also included. |

For the meta-analysis, a study was required to report the following data:

1. the mean RGS score
2. its variance (i.e., SD or SEM)
3. the number of animals per group (N)

We excluded non-rat *in vivo* studies, studies that did not report a control group, and studies that were nor primary research articles (i.e., conference abstracts without data, reviews, book sections, letters, and comments).

### Crowdsourcing and training

Reviewers were individuals who voluntarily participated in this systematic review. All reviewers successfully completed a training module prior to commencing the formal screening and data extraction process. To meet the minimum passing criterion for participating in the screening process, reviewers had to accurately screen 8 out of 10 studies. For participating in the data extraction process, reviewers had to perform extractions for 3 studies, and they were required to correctly identify, and extract information related to the animal characteristics and behavioural outcome measures. A total of 5 independent reviewers took part in data extraction.

### Study selection

All retrieved studies from the systematic search were screened on the Systematic Review Facility (SyRF) platform [31]. Titles and abstracts of studies were screened against the eligibility criteria and then full text screening followed. Screening was conducted by two independent reviewers. A third reviewer reconciled any discrepancies.

### Data extraction

Simultaneously with the full text screening stage on SyRF, two independent reviewers carried out data extraction. Data at the study level were extracted (Table 3), and for studies eligible for meta-analysis, experimental data were extracted (Table 4).

**Table 3.** Study-level data extracted from each included study.

| **Study-level** |  |
| --- | --- |
| **Bibliographic detail** | ▪ First author<br>▪ Year of publication<br>▪ Title |
| **Reporting quality** | Reporting guidelines, such as the ARRIVE (Animal Research: Reporting In Vivo Experiments) [32], which was developed for the purpose to improve the reporting of animal research. The following items were extracted:<br>▪ Reference following a reporting guideline for *in vivo* experimentation (e.g., *ARRIVE*, *Landis4* etc.).<br>▪ Provide evidence of reporting in accordance with the chosen guideline (i.e., *checklist*). |
| **Acclimatisation and animal husbandry** | ▪ Time period of acclimatisation to housing environment following transportation |
| **RGS assessment characteristics** | ▪ Experimental environment (i.e., *light intensity, temperature, noise level, vibration level, light/dark phase, location of the testing environment*) |
| **Curated content** | ▪ Were RGS scores used as an animal welfare measure?<br>▪ Were the authors assessing the sensitivity of the RGS score for this model? |

**Table 4.** Experiment-level data extracted from each included study.

| **Experiment-level** |  |
| --- | --- |
| **Animal** | ▪ Strain<br>▪ Sex<br>▪ Animal supplier<br>▪ Age at the start of experiments<br>▪ Weight |
| **Disease model** | ▪ Control method<br>▪ Method of model induction (surgical, pharmacological, genetic)<br>▪ Perioperative analgesic(s) given before/during/after induction |
| **Intervention** | ▪ Dose<br>▪ Route of administration<br>▪ Co-administration of other drugs<br>▪ Number of administrations<br>▪ Time between drug treatment and model induction |
| **Outcome measure assessment** | Primary outcome – aggregated RGS scores<br>▪ Method of scoring (*i.e., real-time or retrospective scoring, facial units that were scored, scoring scale, how the final score for each facial unit was calculated*)<br>▪ Score reconciliation method (*i.e., number of independent raters, how scores were reconciled*)<br>▪ Calculation of the aggregated score (*i.e., average or sum*)<br>▪ Habituation time<br>▪ Assessment duration<br>▪ Direction of effect<br>▪ Time between the model induction and the first assessment<br>▪ Time between the model induction and the last assessment<br>Secondary outcome – first-ranked stimulus-evoked withdrawal data assessed in the same cohort of animals which participated in the RGS assessment<br>▪ Type of nociceptive assessment and its endpoint<br>▪ Direction of effect |
| **Numerical outcome data** | ▪ Unit<br>▪ Mean outcome<br>▪ Variance<br>▪ Number of animals per group<br>▪ Number of groups served by control group |

The primary outcome of interest was the RGS score. We extracted the aggregated score (i.e., average or sum) of all the facial action units regardless of the scoring method (i.e., real-time vs retrospective scoring).

The secondary outcome of interest was stimulus-evoked withdrawal behaviours assessed in the same cohort of animals that were used in the RGS assessment. Stimulus-evoked withdrawal paradigms were categorised into three types, and the review extracted the first-ranked paradigm type when multiple types are reported:

1. Mechanically evoked (e.g., monofilament evoked limb withdrawal, Randall Selitto)
2. Heat-evoked (e.g., Hargreaves test, hot plate, tail flick)
3. Cold-evoked (e.g., acetone test, cold plate)

This order of ranking was established according to their prevalence in reporting within rodent pain research, arranged from the highest to the lowest. If more than one paradigm type was reported, this information was also annotated.

Graphically presented data were manually extracted using the WebPlotDigitizer. In experiments where the outcome was measured across multiple time points, the time point indicating the maximum effect was extracted. If the reported variance type (i.e., SD or SEM) was unspecified, it was assumed to be SD for a more conservative estimate. The most conservative estimate was utilised when sample size data were presented as a range. When key information (i.e., sample size and variance) was unclear or not reported, attempts were made to contact the corresponding author. If the author did not respond or was unable to provide the necessary information, the study was recorded as having missing data and was excluded from the meta-analysis.

### Risk of bias assessment

A modified version of the CAMARADES checklist and SYRCLE Risk of Bias tool was used to evaluate the reporting of six methodological quality criteria:

1. Random group allocation
2. Allocation concealment
3. Blinding of outcome assessment
4. Sample size calculation
5. Predefined animal inclusion criteria
6. Animal exclusions (i.e., reasons and number of excluded animals)

Reviewers indicated whether each criterion was reported and provided details on the methodology used. Each criterion received an individual rating based on the following criteria:

- Low risk – accepted methods and were adequately described.
- High risk – inappropriate methods that did not efficiently mitigate bias.
- Unclear risk – the methodological quality criterion was not reported, or details of methods were insufficiently reported.

### Other annotations extracted

In addition to the criteria listed above, reviewers extracted statements of potential conflicts of interest, compliance with animal welfare regulations, and adherence to a reporting guideline.

Full annotations can be found in S1 Appendix.

### Data reconciliation

A third independent reviewer compared the data extracted by the initial two independent reviewers, and any disparities were reconciled. In the case of graphically presented data, the SMD effect sizes for each reviewer’s extracted data were calculated for individual comparisons. If the differences in individual comparisons were less than 10%, the reconciler averaged the two means and variance measures. However, if the differences were greater than 10%, the reconciler extracted the outcome data.

### Data analyses

Excluded from the quantitative analysis were studies with insufficient data although their study-level and risk of bias were still evaluated. For studies meeting the eligibility criteria for quantitative analysis, a meta-analysis was performed.

The meta-analysis required a minimum of 10 independent cohort-level comparisons (*k*). If *k* was less than 10, a descriptive summary was presented. Subgroup analyses were undertaken to explore how study characteristics influenced effect sizes. All analyses were performed using R statistical packages: dmetar (version 0.0.9) [33]; meta (version 6.0.0) [34]; and metafor (version 3.8.1) [35].

RGS score data were organised by the type of experiments (i.e., disease modelling or drug intervention experiments). The data was divided into 3 datasets, and they were analysed separately:

i. Disease modelling experiments using internal control groups (i.e., baseline control)
ii. Disease modelling experiments using external control groups (i.e., sham or naïve control)
iii. Drug intervention experiments using vehicle control groups

#### Effect size calculation

For each individual comparison, characterised as a cohort of animals receiving treatment compared to a control group, the effect size was calculated. We planned to calculate the absolute mean difference however, studies did not use the same methods to collect and aggregate the RGS scores (i.e., the same facial action units, summation versus averaging to calculate the aggregated scores). Therefore, effect sizes were calculated using the Hedges’ *g* SMD method. To determine the “true number of control animals”, all sample sizes were adjusted by dividing the reported number of animals in the control group by the number of treatment groups it served. For baseline control experiments, we used equal sample sizes for the control and experimental groups. This was done by dividing the reported sample size by two to prevent double-counting animals in the meta-analysis.

#### Weighting effect sizes

Effect sizes were weighted using the inverse variance method to account for each comparison’s contribution to the overall effect estimate.

#### Summary estimate of effect

Cohort-level effect sizes were aggregated using a random-effects model. The restricted maximum-likelihood method was employed to estimate the variance of the distribution of true effect sizes. Additionally, the Hartung-Knapp-Sidik-Jonkman method was applied to adjust confidence intervals.

#### Heterogeneity

Cochran’s Q and *I^2^* tests were employed to evaluate the presence of heterogeneity, indicating variations in treatment effects observed across different studies. The *P* value for Q was calculated to determine whether all cohort-level comparisons shared a common effect size (*P* > 0.05) or not (*P* < 0.05). The *I^2^* test quantified the proportion of total variance between studies attributed to true differences in effect sizes rather than chance. *I^2^* values were interpreted following the definitions provided by Higgins and Thompsons [36]:

- to 25% indicates very low heterogeneity.
- to 50% indicates low heterogeneity.
- to 75% indicates moderate heterogeneity.
- >75% indicates high heterogeneity.

#### Subgroup analyses

To explore potential sources of heterogeneity, stratified subgroup analyses for categorical variables were performed according to:

- Strain
- Sex
- Model class
- Drug class
- Type of pain (i.e., acute nociception or persistent pain-related disease conditions)
- Items of study quality criteria

Multivariate meta-regressions were planned to identify other factors relating to experimental conditions and RGS assessment characteristics that could influence the RGS scores (Table 5).

**Table 5.** Experimental design variables relating experimental conditions and RGS assessment characteristics.

|  |  |
| --- | --- |
| **Experimental conditions** | ▪ Light intensity of the testing room<br>▪ Temperature of the testing room<br>▪ Noise level of the testing room<br>▪ Vibration level of the testing room<br>▪ RGS assessment conducted in the light or dark phase |
| **RGS assessment characteristics** | ▪ Number of raters<br>▪ Rater agreement analysis |
|  | ▪ RGS score reconciliation<br>▪ Method of RGS scoring<br>▪ Method of RGS calculation<br>▪ Facial units that were scored<br>▪ Habituation time<br>▪ Time between model induction and the first RGS assessment<br>▪ Time between model induction and the last RGS assessment |

#### Multilevel meta-analyses

To assess the heterogeneity and relationship between experimental variables within the model, a post-hoc multilevel meta-analysis was conducted.

#### Publication bias

Funnel plots were plotted to visually examine asymmetry in the plot. SMDs were plotted against a precision estimate based on sample size (1/√N). Egger’s regression test was used to statistically assess the funnel plot asymmetry. Additionally, a Trim-and-Fill analysis was conducted to address any funnel plot asymmetry. This involved imputing theoretically missing studies, allowing for a recalculation of the effect size.

#### Correlation of thigmotaxis and stimulus-evoked limb withdrawal behavioural outcomes

Cohort-level comparisons of experiments that assessed both RGS scores and stimulus-evoked limb withdrawal were used to investigate correlation. A Pearson’s Correlation test was used.

#### Sensitivity test

Outliers were excluded to investigate whether a single study or group of studies have skewed the analysis.

## Results

### Study selection

A total of 162 publications were retrieved from the systematic searches. After title and abstract screening, 89 studies were included. Full text screening led to the inclusion of 54 studies, and their data were extracted for the meta-analysis (Fig 1).

**Fig 1.**
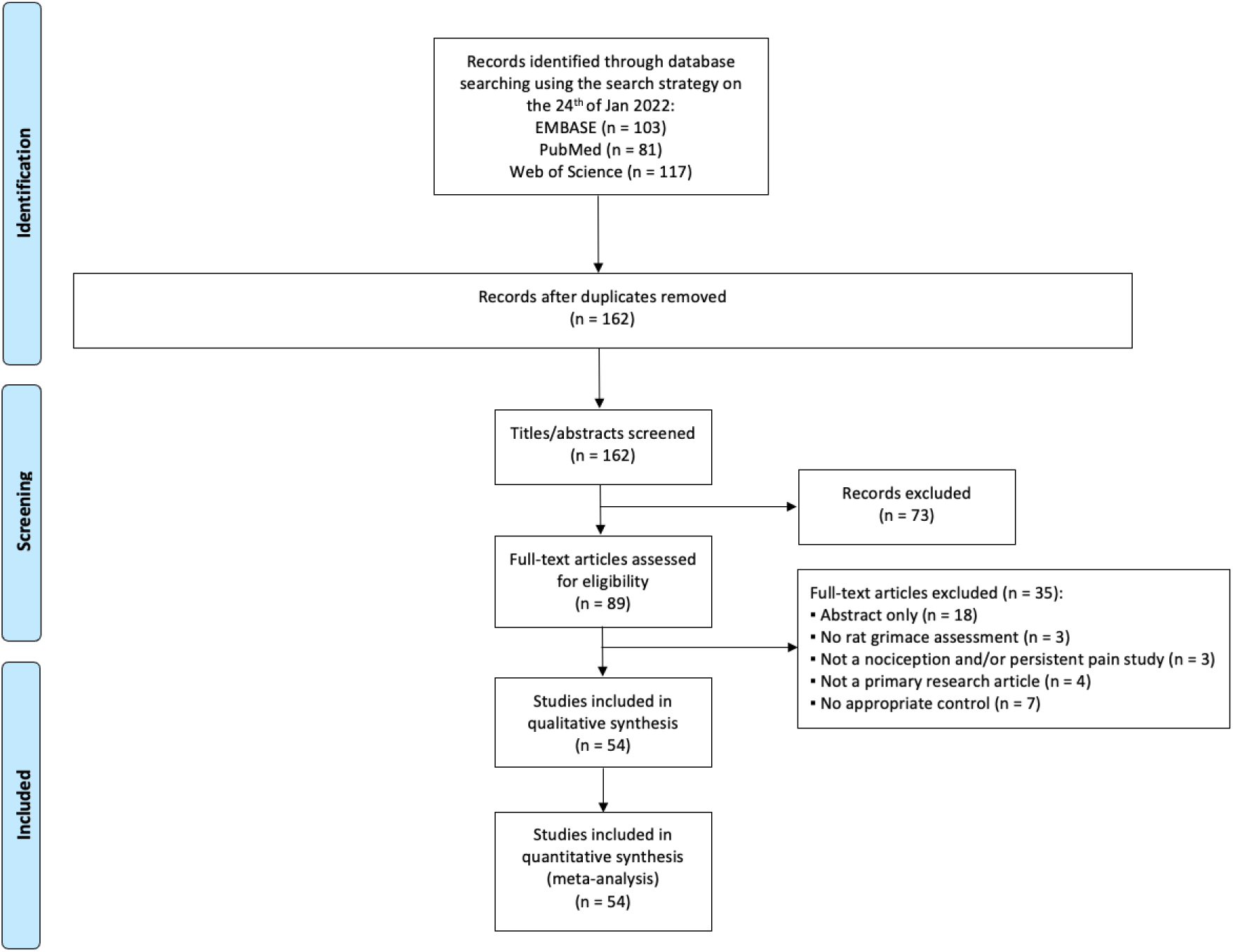
A flow diagram of RGS systematic review. Publications identified through the systematic searches of three electronic databases (EMBASE, PubMed, and Web of Science). The diagram illustrates the number of records (n) at deduplication, screening, and study eligibility for both qualitative and quantitative analyses. Reported in accordance with the PRISMA 2020 guidelines [37].

#### Study characteristics

The 54 studies included a total of 1883 rats (849 in disease modelling and 1034 in drug intervention experiments).

RGS scores were reported in 27 different rat disease models associated with persistent pain. These models are listed according to the classification (15 model classes) in Table 6; post-surgical (39%, *k* = 54), and orofacial inflammation (19%, *k* = 26) were the most frequently reported. No studies assessed the effects of acute nociception assays on the RGS scores.

**Table 6.** Summary of the model classes used in disease modelling and drug intervention experiments within the RGS systematic review.

| Model Class | Model Name | Disease Modelling Experiments |  |  |
| --- | --- | --- | --- | --- |
|  |  | No. of studies | No. of *k* | No. of rats |
| Post-surgical | Laparotomy | 10 | 40 | 354 |
|  | Paw surgical incision | 3 | 11 | 129 |
|  | Craniotomy | 1 | 2 | 32 |
|  | Stereotaxic surgery induced tympanic membrane rupture | 1 | 1 | 7 |
| Orofacial inflammation | Orthodontic tooth movement induced inflammation | 7 | 16 | 502 |
|  | Temporomandibular joint mechanical loading | 3 | 6 | 56 |
|  | Temporomandibular joint osteoarthritis | 1 | 3 | 24 |
|  | Occlusal dental interference | 1 | 1 | 60 |
| Neuropathy – nerve injury | Infraorbital nerve constriction | 3 | 3 | 74 |
|  | Sciatic nerve constriction | 1 | 1 | 16 |
|  | Compression of cervical nerve root | 1 | 1 | 14 |
| Inflammation | CFA (non-articular) | 3 | 7 | 71 |
|  | Carrageenan | 2 | 5 | 80 |
|  | LPS | 1 | 1 | 11 |
| Burn injury | Second-degree burn | 2 | 2 | 40 |
| Traumatic brain injury | Cranium Only Blast Injury | 2 | 2 | 47 |
| Reserpine | Reserpine | 1 | 11 | 106 |
| Combined: inflammation +<br>Post-surgical | LPS + paw surgical incision | 1 | 9 | 91 |
| Spinal cord injury | Laminectomy | 1 | 4 | 24 |
|  | Spinal cord injury | 1 | 1 | 15 |
| Stroke | Collagenase-induced intracerebral haemorrhage | 1 | 3 | 33 |
| Visceral inflammation | DSS induced colitis | 1 | 2 | 41 |
| Meningeal neurogenic inflammation | Chemical stimulation of meninges | 1 | 2 | 11 |
|  | Nitroglycerine | 1 | 1 | 18 |
| Nerve growth factor induced inflammation of the L5 vertebrae | Nerve growth factor | 1 | 1 | 12 |
| Formalin | Formalin | 1 | 1 | 9 |
| Arthropathy | Carrageenan/Kaolin | 1 | 1 | 6 |
| **Total** |  |  | 138 | 1883 |

The total number of studies is not provided as summation will surpass the true total (54 studies) because of multiple disease models being investigated per study. $k$ , independent cohort-level effect.

Twenty-seven drugs are classified by their mechanism of action (16 drug classes) using the IUPHAR/BPS guide to Pharmacology as listed in Table 7; NSAIDs (33%, *k* = 29) were the most frequently investigated.

**Table 7.**
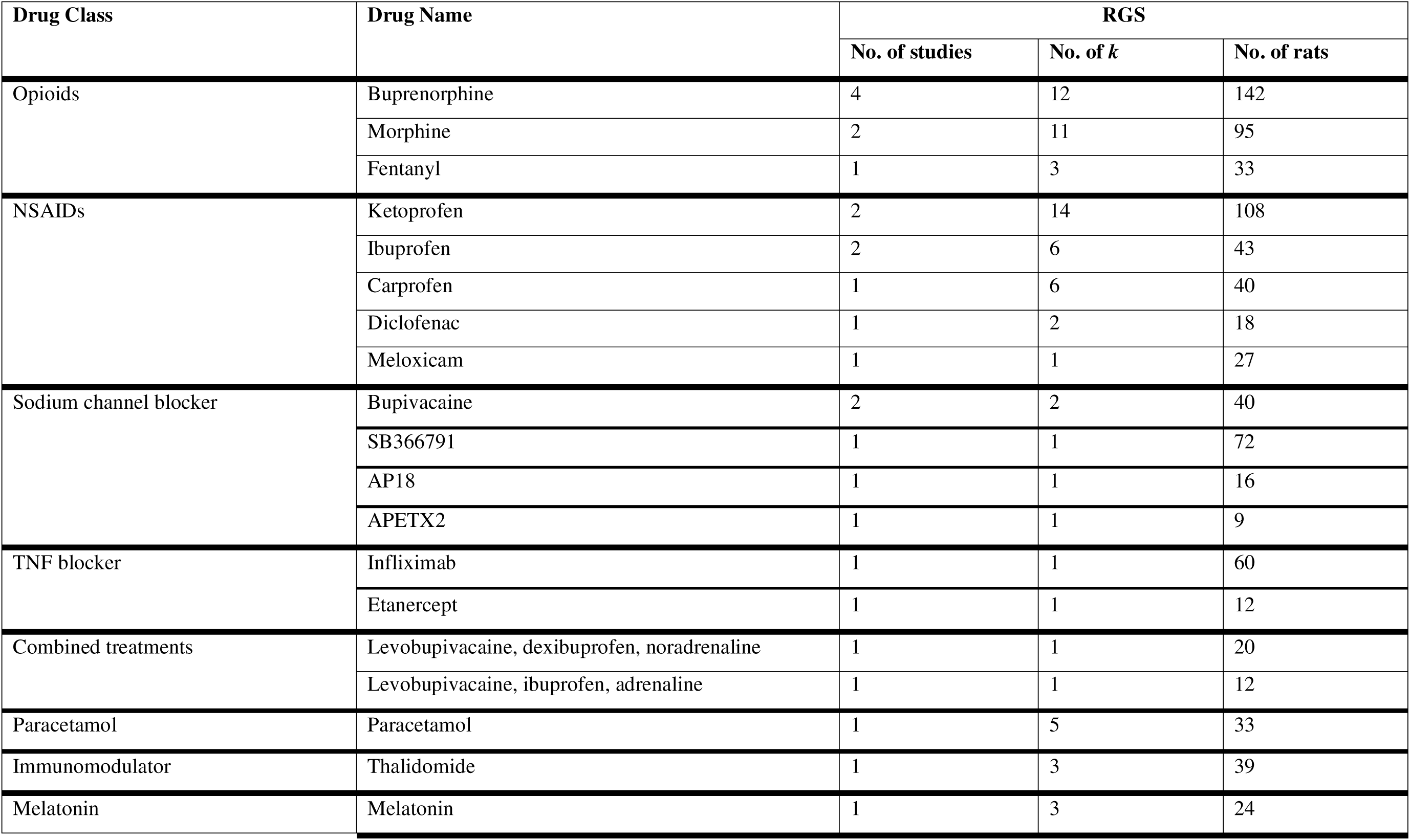
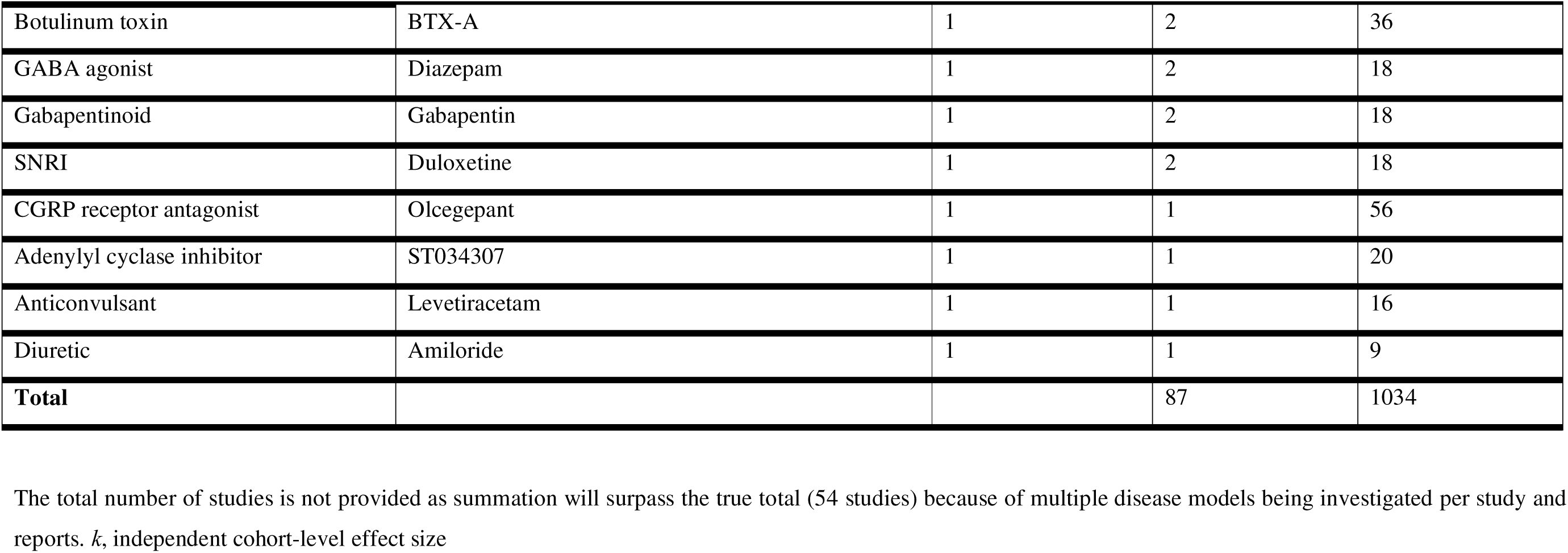
Summary of the drug classes used to assess the effect on RGS scores in rat disease models associated with persistent pain.

#### Meta-analysis of RGS scores

For disease modelling experiments, two types of controls were extracted:

- External control (i.e., a separate cohort of sham or naïve animals)
- Internal control (i.e., a baseline grimace score)

There were fewer comparisons using internal baseline controls:

- External control (*k* = 36)
- Internal control (*k* = 15)

Based on the differences between the two control types, they were analysed separately. Due to the small number of individual cohort-level experimental comparisons involving internal baseline control data, the manuscript prioritizes the presentation and interpretation of disease modelling experiments utilizing external controls. S2 Appendix contains detailed information on data characteristics and the meta-analysis of internal baseline control data.

#### Disease modelling experiments

A total of 26 studies, encompassing 36 cohort-level comparisons, 709 rats, assessed the effects of 12 classes of disease models associated with persistent pain on RGS scores (Table 8). The sample size ranged from 7 to 73 animals per group, with a median of 16. Orofacial inflammation was the most reported model class accounting for 31 % of studies (*k* = 11). No studies included acute nociception assays. Among rat strains, the Sprague Dawley was used most often (53%, *k* = 19). Male rats were used in 83% (*k* = 30) of the experiments, whereas females were used in 8% (*k* = 3), and 8% (*k* = 3) reported mixed sex groups.

**Table 8.** The number of cohort-level comparisons and animals for animal model characteristics used in RGS disease modelling experiments using external controls.

| | No. of studies | No. of $k$ | No. of animals |
| --- | --- | --- | --- |
| **Disease Model** |  |  |  |
| Orofacial inflammation | 8 | 11 | 324 |
| Post-surgical | 7 | 8 | 110 |
| Inflammation | 3 | 4 | 57 |
| Neuropathy - nerve injury | 3 | 3 | 49 |
| Traumatic brain injury | 2 | 2 | 47 |
| Visceral inflammation | 1 | 2 | 41 |
| Meningeal neurogenic inflammation | 1 | 1 | 18 |
| Reserpine | 1 | 1 | 16 |
| Spinal cord injury | 1 | 1 | 15 |
| Nerve growth factor induced inflammation of the L5 vertebrae | 1 | 1 | 12 |
| Inflammation + Post-surgical | 1 | 1 | 11 |
| Formalin | 1 | 1 | 9 |
| **Rat Strain** |  |  |  |
| Sprague Dawley | 15 | 19 | 450 |
| Wistar | 11 | 16 | 245 |
| Holtzman | 1 | 1 | 14 |

| Sex |  |  |  |
| --- | --- | --- | --- |
| Male | 23 | 30 | 625 |
| Mixed | 2 | 3 | 48 |
| Female | 2 | 3 | 36 |

#### RGS scores were increased by disease models associated with persistent pain

Disease models associated with persistent pain significantly worsened RGS scores in experiments using external controls, a summary effect size of SMD = −4.44 [95%CI −6.25 to −2.63]). Heterogeneity was high (*Q* = 6916.58, *df* = 35, *P* < 0.0001, *I^2^* = 99.5%) (Fig 2). Sensitivity analysis showed that the removal of the three outlying comparisons from Guo et al. (2019), Gao et al. (2016), and Long et al. (2015) which reported very large effect sizes, reduced the summary effect size to – 2.96 [95%CI −3.57 to −2.35].

**Fig 2.**
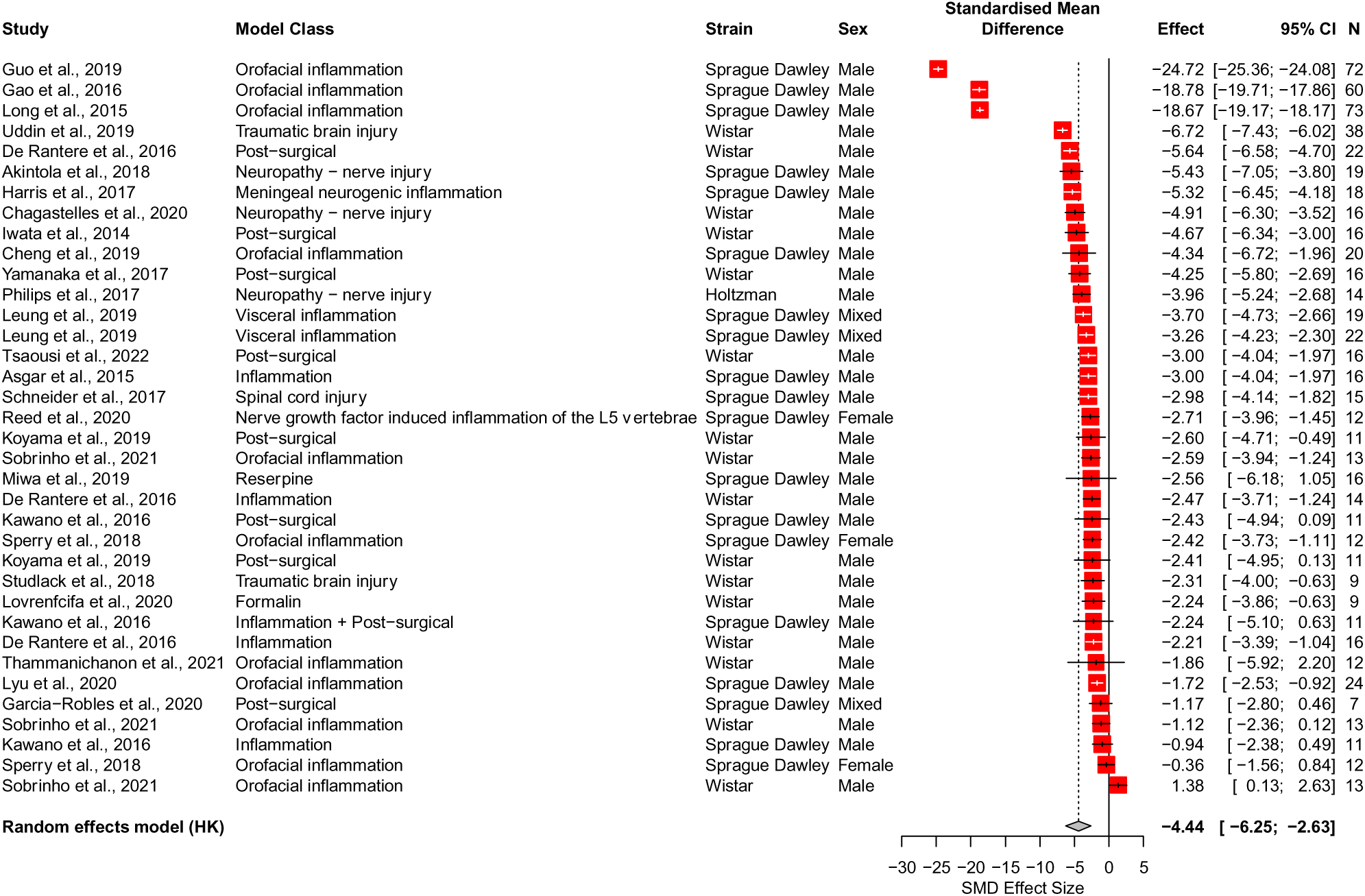
Effects of disease models on RGS scores: summary forest plot. A summary forest plot of the 36 cohort-level comparisons which assessed the impact of disease modelling on RGS scores using a separate group of control rats. For each comparison, an effect size was calculated using the Hedges’ g SMD method. Effect sizes were pooled using the random effects model. The restricted maximum-likelihood method was used to estimate heterogeneity. The overall effect size is −4.44 [95% CI −6.25 to −2.63]; *Q* = 6916.58, *df* = 35, *P* < 0.0001, *I^2^* = 99.5%. The size of the square represents the weight, which reflects the contribution of each comparison with the pooled effect estimate. CI, confidence interval; N, number of animals.

#### Strain did not influence RGS scores

Three rat strains were reported. Sprague Dawley, the most commonly used strain (53%, *k* = 19) and Wistar (43%, *k* = 15) were the only two strains with sufficient data for stratified subgroup analysis. Strain did not account for a significant proportion of heterogeneity (Q = 2.10, *df* = 1, *P* = 0.15) (Fig 3).

**Fig 3.**
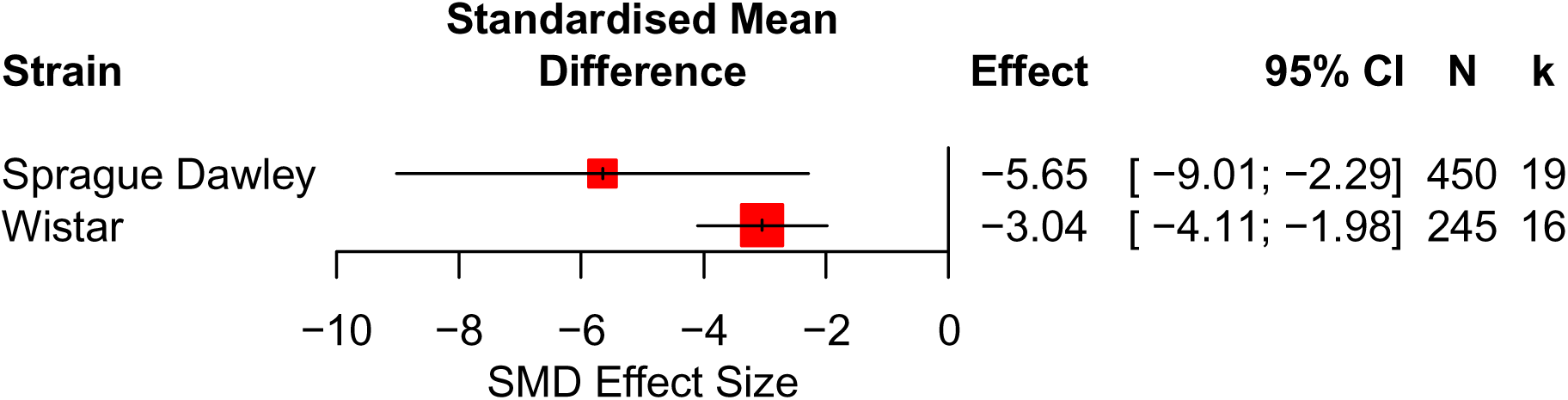
Stratified subgroup analyses of RGS scores. A forest plot of the RGS scores stratified by rat strains in rats modelled with diseases associated with persistent. The size of the square represents the weight. CI, confidence interval; *k*, number of cohort-level comparisons; N, number of animals.

The effects of model type, sex and other experimental factors could not be analysed due to insufficient reporting (k < 10 for each characteristic). A summary of these characteristics is provided in Section 3.4. Reporting of other study characteristics and experimental factors.

#### Drug intervention experiments

A total of 23 studies, containing 87 cohort-level comparisons, 1034 rats, investigated the effects of drug treatments from 16 drug classes on RGS across seven models associated with persistent pain (Table 9). Sample sizes ranged from 4 to 72 animals per group, with a median of 9. Post-surgical models were the most reported (51%, *k* = 44) and opioids the most commonly studied drug (30%, *k* = 26). No studies examined drug treatment effects in acute nociception assays. There were 4 different rat strains reported, and Sprague Dawley was the most commonly used (51%, *k* = 44). Male rats predominated in 51% (*k* = 44) of experiments, 41% (*k* = 36) used mixed sexes, 7% (*k* = 6) used female animals, and 1% (*k* = 1) did not report the sex.

**Table 9.** The number of cohort-level comparisons and animals for animal model characteristics used in RGS drug intervention experiments.

|  | No. of studies | No. of *k* | No. of animals |
| --- | --- | --- | --- |
| **Disease Model** |  |  |  |
| Post-surgical | 8 | 44 | 398 |
| Orofacial inflammation | 8 | 12 | 298 |
| Inflammation | 3 | 8 | 95 |
| Burn injury | 2 | 2 | 40 |
| Reserpine | 1 | 10 | 90 |
| Inflammation + Post-surgical | 1 | 8 | 80 |
| Stroke | 1 | 3 | 33 |
| **Drug Class** |  |  |  |
| Opioid | 7 | 26 | 270 |
| NSAID | 5 | 29 | 236 |
| Sodium channel blocker | 5 | 5 | 137 |
| TNF blocker | 2 | 2 | 72 |
| Paracetamol | 1 | 5 | 33 |
| Immunomodulator | 1 | 3 | 39 |
| Melatonin | 1 | 3 | 24 |
| BTX-A | 1 | 2 | 36 |
| GABA agonist | 1 | 2 | 18 |
| Gabapentinoid | 1 | 2 | 18 |
| SNRI | 1 | 2 | 18 |
| CGRP receptor antagonist | 1 | 1 | 56 |
| Adenylyl cyclase inhibitor | 1 | 1 | 20 |
| Levobupivacaine + Dexibuprofen + Norepinephrine | 1 | 1 | 20 |
| Anticonvulsant | 1 | 1 | 16 |
| Levobupivacaine + ibuprofen + Epinephrine | 1 | 1 | 12 |
| Diuretic | 1 | 1 | 9 |
| **Rat Strain** |  |  |  |
| Sprague Dawley | 12 | 44 | 589 |
| Wistar | 7 | 37 | 336 |
| Not reported | 3 | 4 | 77 |
| Wister albino | 1 | 1 | 20 |
| Holtzman | 1 | 1 | 12 |
| **Sex** |  |  |  |
| Male | 12 | 44 | 580 |
| Mixed | 5 | 36 | 269 |
| Female | 5 | 6 | 165 |
| Not reported | 1 | 1 | 20 |

#### Drug treatments improved RGS scores in models of persistent pain

Overall, drug treatments significantly improved RGS scores in disease models associated with persistent pain, with a summary effect size of SMD = 1.97 [95%CI 1.25 to 2.69]. Heterogeneity was high (*Q* = 5474.43, *df* = 86, *P* < 0.0001, *I^2^* = 98.4%) (Fig 4). Sensitivity analysis showed that the removal of the three comparisons from Long et al. (2015) and Guo et al. (2019) which reported very large effect sizes, reduced the summary effect size to 1.64 [95%CI 1.25 to 2.03].

**Fig 4.**
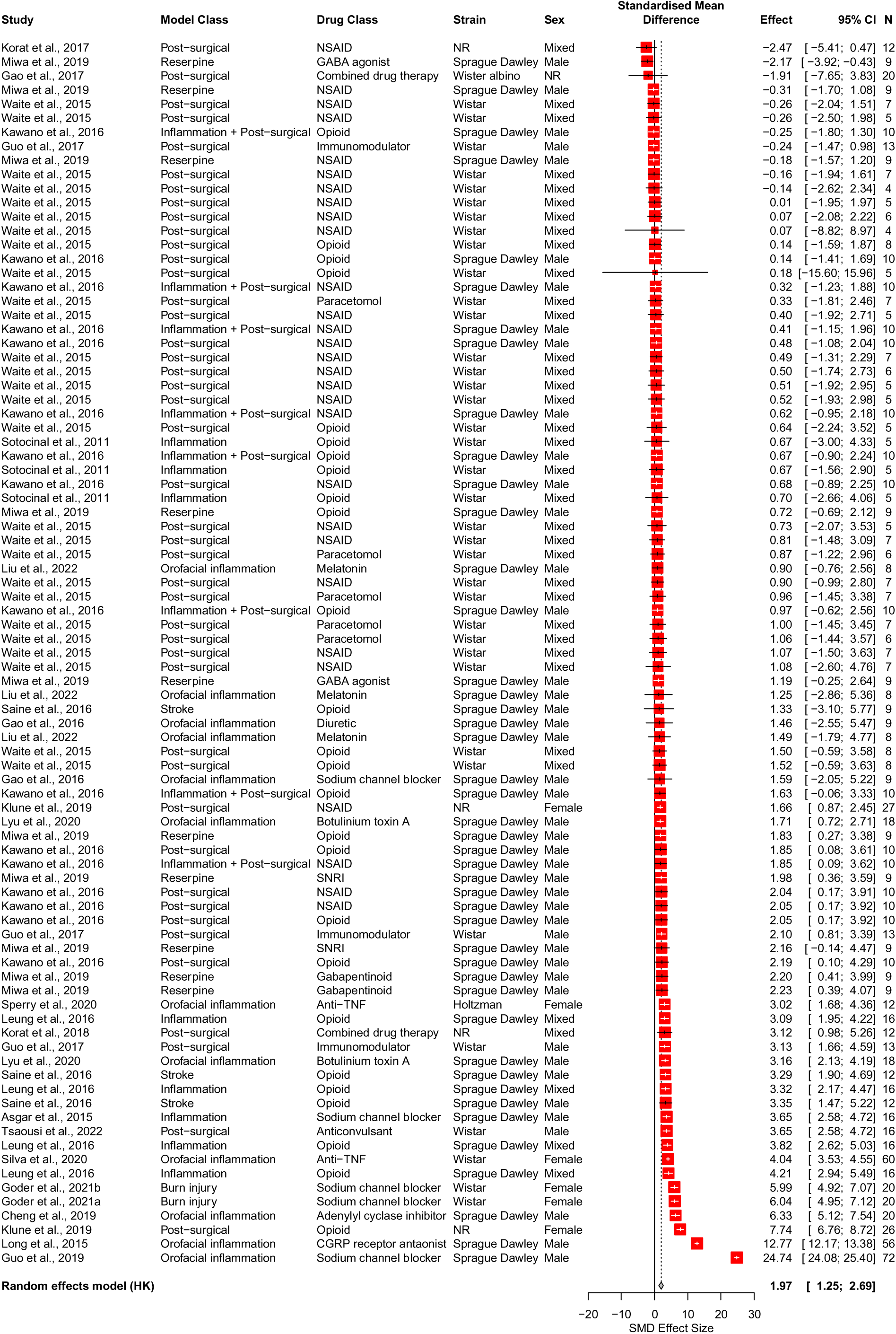
Effects of drug treatments on RGS scores: summary forest plot. A summary forest plot of the 87 cohort-level comparisons which assessed the effects of drug treatments on RGS scores in rats modelled with experimental pain. For each comparison, an effect size was calculated using the Hedges’ g SMD method. Effect sizes were pooled using the random effects model. The restricted maximum-likelihood method was used to estimate heterogeneity. The overall effect size is 1.97 [95% CI 1.25 to 2.69]; *Q* = 5474.43, *df* = 86, *P* < 0.0001, *I^2^* = 98.4%. The size of the square represents the weight, which reflects the contribution of each comparison with the pooled effect estimate. CI, confidence interval; N, number of animals; NR, not reported.

Across the studies, heterogeneity was partially explained by differences in animal and drug characteristics. Seven model classes were reported. Post-surgical was the most reported (51%, *k* = 44). Post-surgical, orofacial inflammation and reserpine provided sufficient data for analysis. However, the model class did not account for a significant proportion of heterogeneity (Q = 3.51, *df* = 2, *P* = 0.17) (Fig 5A).

**Fig 5.**
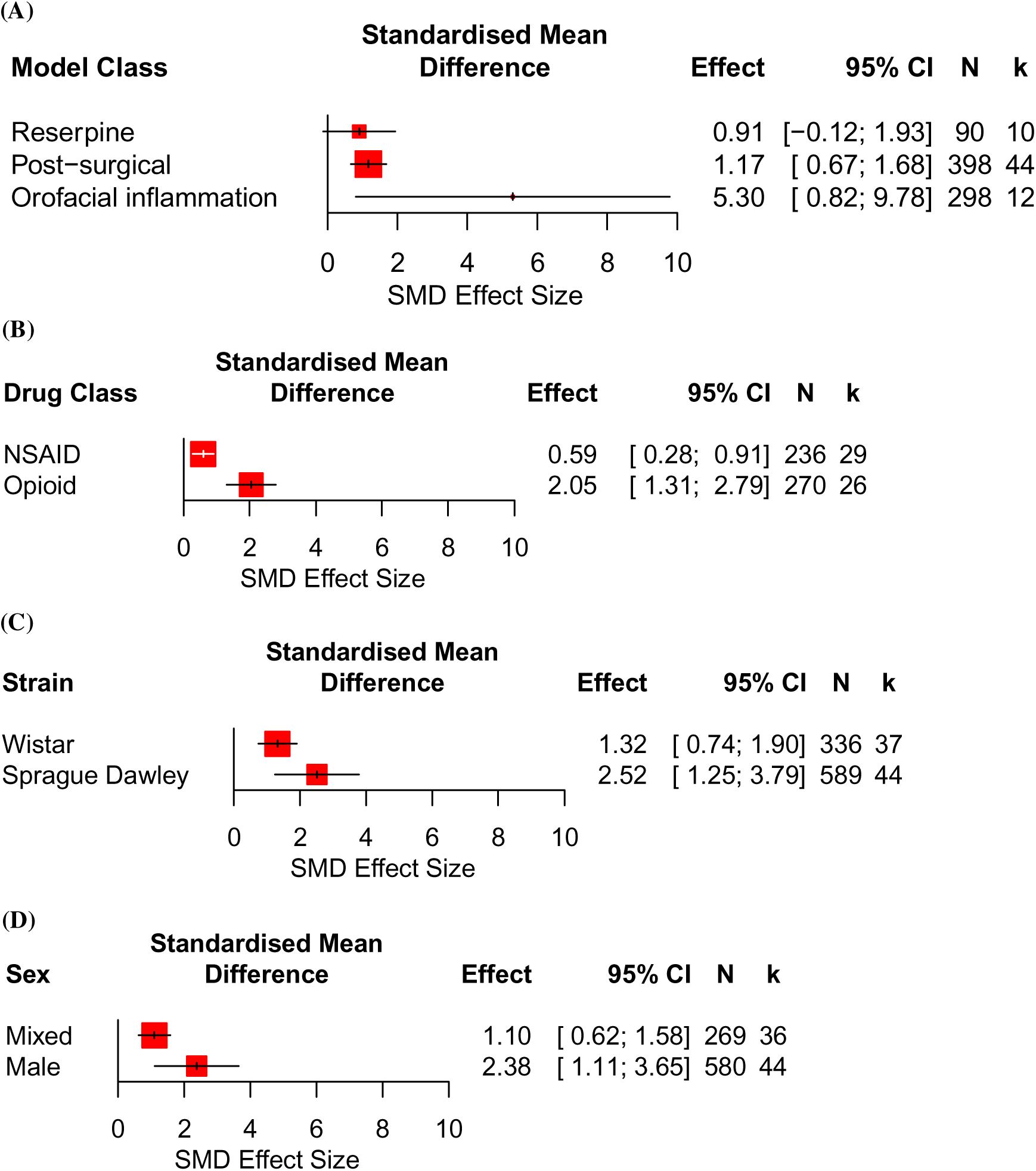
Stratified subgroup analyses of RGS scores in drug treated rats. Forest plots of the RGS scores stratified by: (A) model class, (B) drug class, (C) strain, and (D) sex. The size of the square represents the weight. CI, confidence interval; *k*, number of cohort-level comparisons; N, number of animals.

Drug class contributed to variability. Sixteen drug classes were reported. NSAIDs were the most studied (33%, *k* = 29) followed by opioids (30%, *k* = 26). NSAIDs and opioids had enough data for stratified subgroup analysis, which revealed that drug class accounted for a significant proportion of heterogeneity (Q = 12.47, *df* = 1, *P* = 0.0004) (Fig 5B).

Other factors, such as strain and sex had less influence on heterogeneity. Sprague Dawley and Wistar rats were the only strains with sufficient data for subgroup analysis but strain did not account for a significant proportion of heterogeneity (Q = 2.83, *df* = 1, *P* = 0.09) (Fig 5C). Similarly, while male rats showed the largest reductions in RGS scores following drug treatments, sex did not account for a significant proportion of heterogeneity (Q = 3.42, *df* = 1, *P* = 0.06) (Fig 5D).

A multilevel meta-analysis further explored sources of heterogeneity, using a six-level clustering model based on hierarchical factors such as model class, drug class, strain, sex and other experimental variables. At the highest level, level 6 was defined by the model class, and was followed by level 5 for the drug class, level 4 for the strain, level 3 for the sex, level 2 for the other undefined experimental factors, and level 1 for the within-study variance. This post-hoc analysis did not enforce the k >10 requirement for inclusion.

The analysis showed that sampling error variance at level 1 was low (0.58, *I^2^* = 3.59%) and the overall between-study heterogeneity was high (*I^2^* = 96.41%) (Fig 6). Amongst the defined factors, 35.14% of the heterogeneity was explained by drug class difference 18.65% was explained by model class difference, 16% was attributed to sex, 0.01% was explained by strain difference, while the remaining 26.62% could not be explained by the variables that were defined in this model.

**Fig 6.**
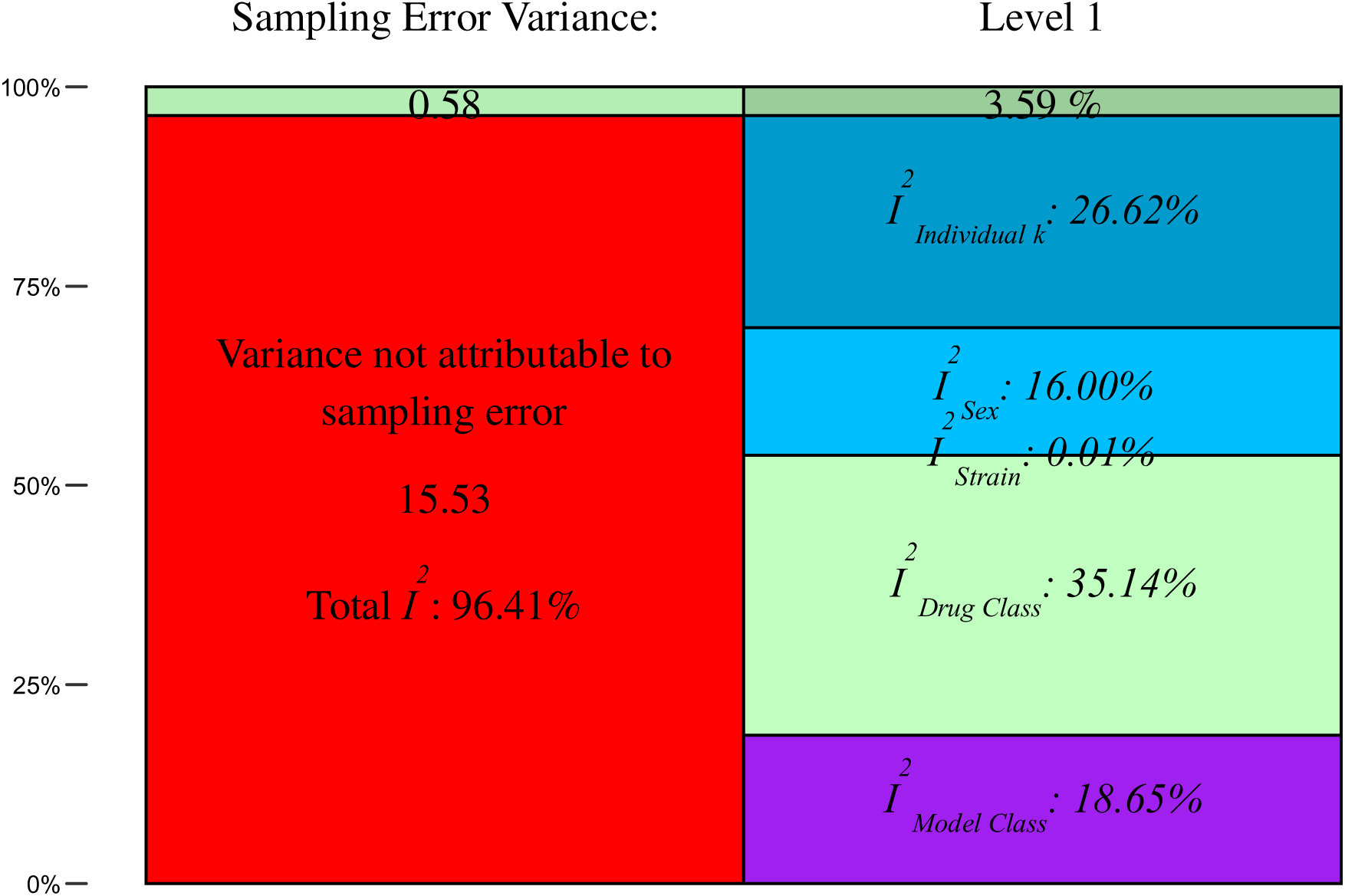
Multilevel meta-analysis of RGS scores in drug treated rats. A plot for the distribution of the total variance in the seven-level model analysis (with each level defined as follows: level 6 by the model class, level 5 by the drug class, level 4 by the strain, level 3 by the sex, level 2 by the undefined experimental factors, and level 1 by the within-study variance). The overall heterogeneity was high, as indicated by an *I^2^* value of 96.41%. The choice of drug class explained the most amount of heterogeneity. 30.01% of heterogeneity could not be explained by the variables defined in this model. A random-effects model was used for pooling the cohort-level effect sizes. *t* test was used to apply the regression coefficients. The restricted maximum-likelihood method was used to estimate the variance of the distribution of true effect sizes.

#### Effects of drug class on the RGS scores in rats

A *post-hoc* stratified meta-analysis evaluated the association between study characteristics and RGS scores in rats treated by the two most frequently reported drug classes: NSAIDs and opioids. It is important to note that the usual criterion of requiring a minimum of 10 independent cohort-level effect sizes (*k*) for each variable was not applied in this analysis.

#### NSAIDs

Ketoprofen (SMD = 0.77 [95% CI 0.35 to 1.19]) significantly reduced the RGS scores in rats modelled with persistent pain (Fig 7A). However, for other NSAIDs, the number of cohort-level comparisons was below 10. NSAIDs were assessed in 3 model classes, with post-surgical models – the only class meeting the *k* >10 threshold – showing significant improvement in RGS score (Fig 7B). Analysis by rat strain revealed significant improvement in RGS scores for both Sprague Dawley and Wistar strains, although two cohort-level comparisons did not specify strain (Fig 7C). Male rats treated with NSAIDs showed significant reductions in RGS scores, while data for females were limited to a single cohort-level comparison (Fig 7D).

**Fig 7.**
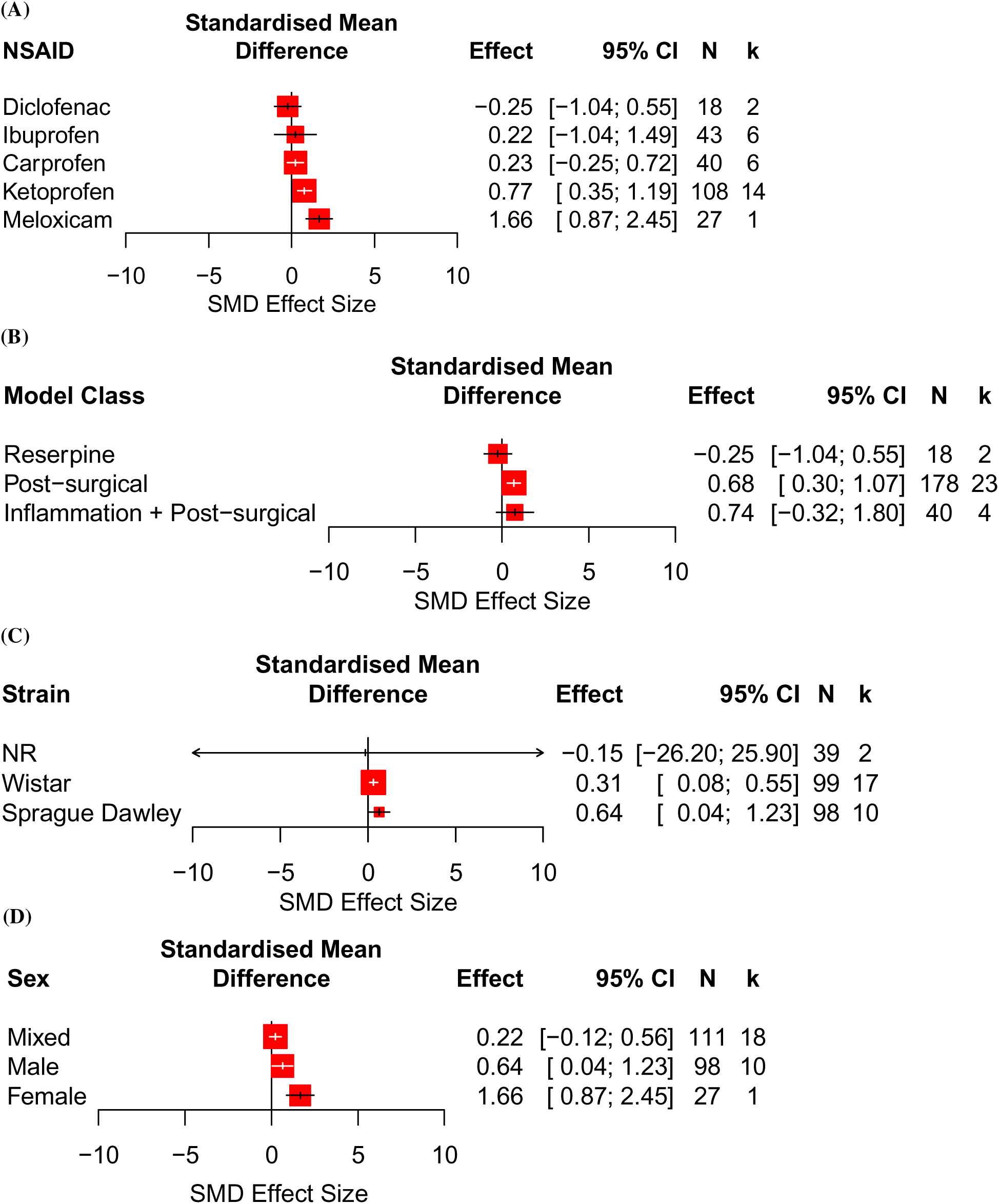
Stratified subgroup analyses of RGS scores in rats treated with NSAIDs. Forest plots of treatment effects of NSAIDs on RGS scores in rats modelled with persistent pain: (A) type of NSAID, (B) model class, (C) strain, and (D) sex. The size of the square represents the weight. CI, confidence interval; *k*, number of cohort-level comparisons; N, number of animals; SMD, standardised mean difference.

#### Opioids

Morphine and buprenorphine significantly improved RGS scores in rats modelled with persistent pain (Fig 8A). Opioids were evaluated in 5 model classes, with post-surgical models – the only class meeting the *k* >10 threshold – showing significant improvement (Fig 8B). By strain, significant improvements in RGS were observed for Sprague Dawley rats, while one cohort level comparison did not report the strain (Fig 8C). Opioids significantly improved RGS scores in experiments conducted in male and mixed sexes experimental groups, with a larger treatment effect observed in mixed-sex groups. Data for females were limited to a single cohort-level comparison (Fig 8D).

**Fig 8.**
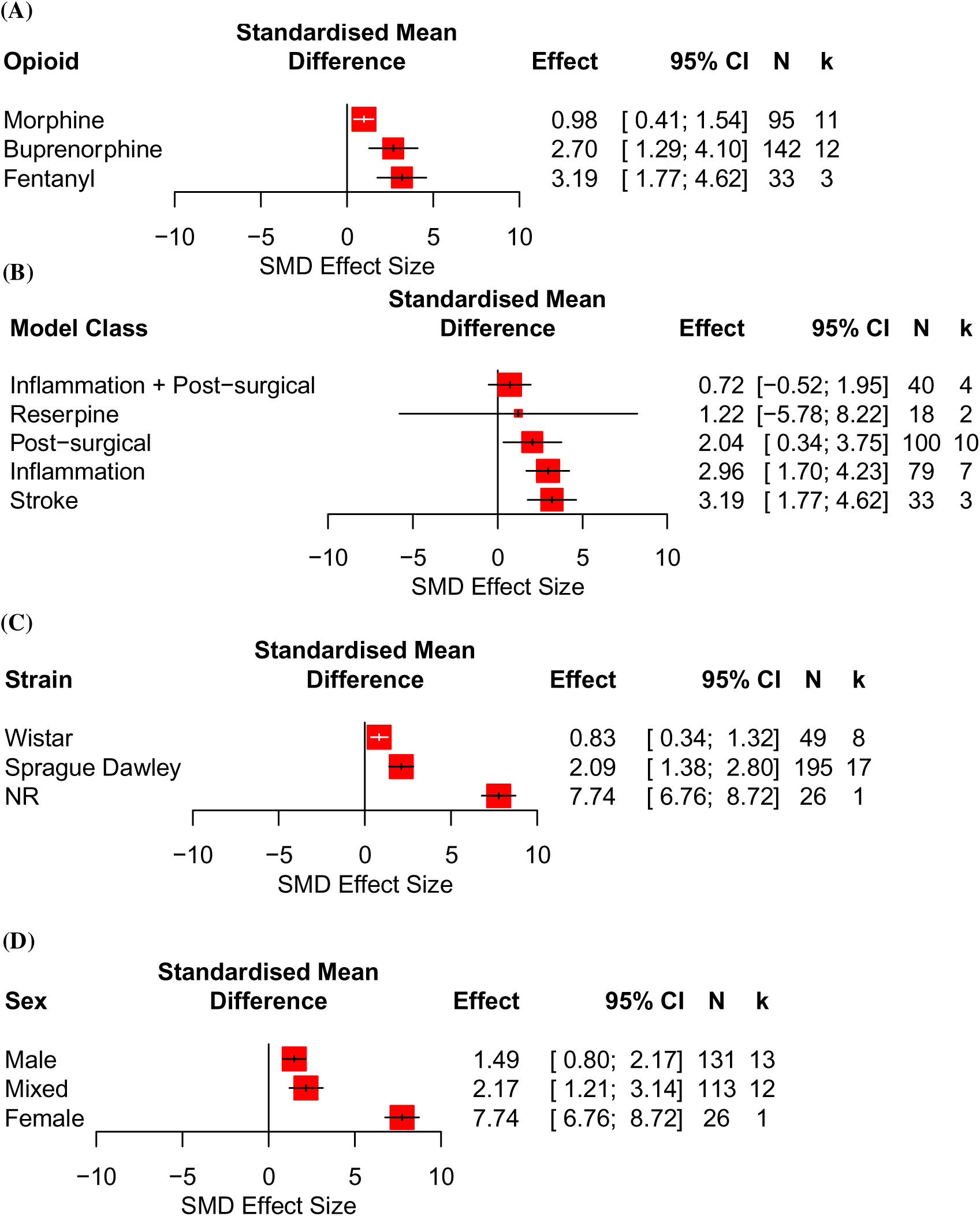
Stratified subgroup analyses of RGS scores in rats treated with opioids. Forest plots of treatment effects of opioids on RGS scores in rats modelled with persistent pain: (A) type of opioid, (B) model class, (C) strain, and (D) sex. The size of the square represents the weight. CI, confidence interval; *k*, number of cohort-level comparisons; N, number of animals; SMD, standardised mean difference.

Information regarding experimental conditions, RGS assessment characteristics and animal cohort characteristics for each study are summarised in Table 10.

**Table 10.** Number of studies which reported relevant information on experimental conditions, RGS assessment characteristics and animal cohort characteristics.

| Item reported | Number of studies | % | Data |
| --- | --- | --- | --- |
| Acclimatisation period before experiments | 16 | 30 | Range = 48 to 168 hr<br>Median = 168 hr |
| Light intensity in the testing room | 0 | 0 | - |
| Temperature in the testing room | 1 | 2 | 23°C = 1 study |
| Vibration level in the testing room | 0 | 0 | - |
| Noise level in the testing room | 3 | 6 | 45dB = 1 study<br><45dB = 2 studies |
| In which phase, was the RGS assessed | 8 | 15 | Light = 8 studies |
| Location of the testing environment | 36 | 67 | Familiar home cage = 1 study<br>Novel testing cage = 35 studies |
| Authors reported rater training for the RGS scoring | 6 | 11 | Yes = 6 studies |
| RGS score was rated by | 34 | 63 | Single rater = 24 studies<br>Multiple raters = 10 studies |
| RGS score was rated independently | 7 | 13 | Yes = 7 studies |
| A rater agreement analysis conducted | 5 | 9 | Yes = 5 studies |
| Score reconciliation method | 6 | 11 | Average the scores = 6 studies |
| Method of RGS scoring | 40 | 74 | Retrospective image = 38 studies<br>Real-time = 2 studies |
| Method of frame selection for RGS scoring | 33 | 61 | Manual selection = 26 studies<br>Automated selection tool = 7 studies |
| Conventional facial units scored | 47 | 87 | Yes = 40 studies<br>No = 7 studies |
| Method of calculating the final aggregated score | 35 | 65 | Average = 31 studies<br>Sum = 4 studies |
| Habituation test to the RGS assessment | 19 | 35 | Range = 5 to 2880 minutes<br>Median = 10 minutes |
| Assessment duration | 33 | 61 | Range = 0.03 to 30 hr<br>Median = 0.5 hr |
| Time between model induction and the first RGS assessment | 45 | 83 | Range = 0 to 840 hr<br>Median = 2 hr |
| Time between model induction and the last RGS assessment | 45 | 83 | Range = 0 to 804 hr<br>Median = 144 hr |
| Animal supplier | 34 | 63 | Most reported = Charles River |
| Age | 25 | 46 | Range = 35 to 735 days<br>Median = 60 days |
| Weight | 41 | 76 | Range = 157.5 to 595 g |
|  |  |  | Median = 275 g |
| N number | 54 | 100 | Range = 4 to 73 animals<br>Median = 10 animals |

**3.5. Correlation of RGS and mechanically evoked limb withdrawal outcomes**. Stimulus-evoked limb withdrawal data were extracted based on a hierarchy system. Mechanically evoked limb withdrawal was the first ranked stimulus-evoked outcome data. The included studies that also investigated stimulus-evoked behavioural outcomes all reported mechanically evoked limb withdrawal outcomes. Correlations could not be assessed for heat and cold evoked limb withdrawal outcomes due to insufficient data. The number of studies and cohort-level comparisons are listed in Table 11.

**Table 11.** The number of studies that investigated RGS scores and mechanically induced limb withdrawal outcomes.

| Experiment type | No. of studies | No. of cohort-level comparisons (*k*) |
| --- | --- | --- |
| RGS disease modelling | 9 | 14 |
| RGS drug intervention | 3 | 14 |

The reported model classes for disease modelling and drug intervention experiments can be found in Table 12.

**Table 12.**
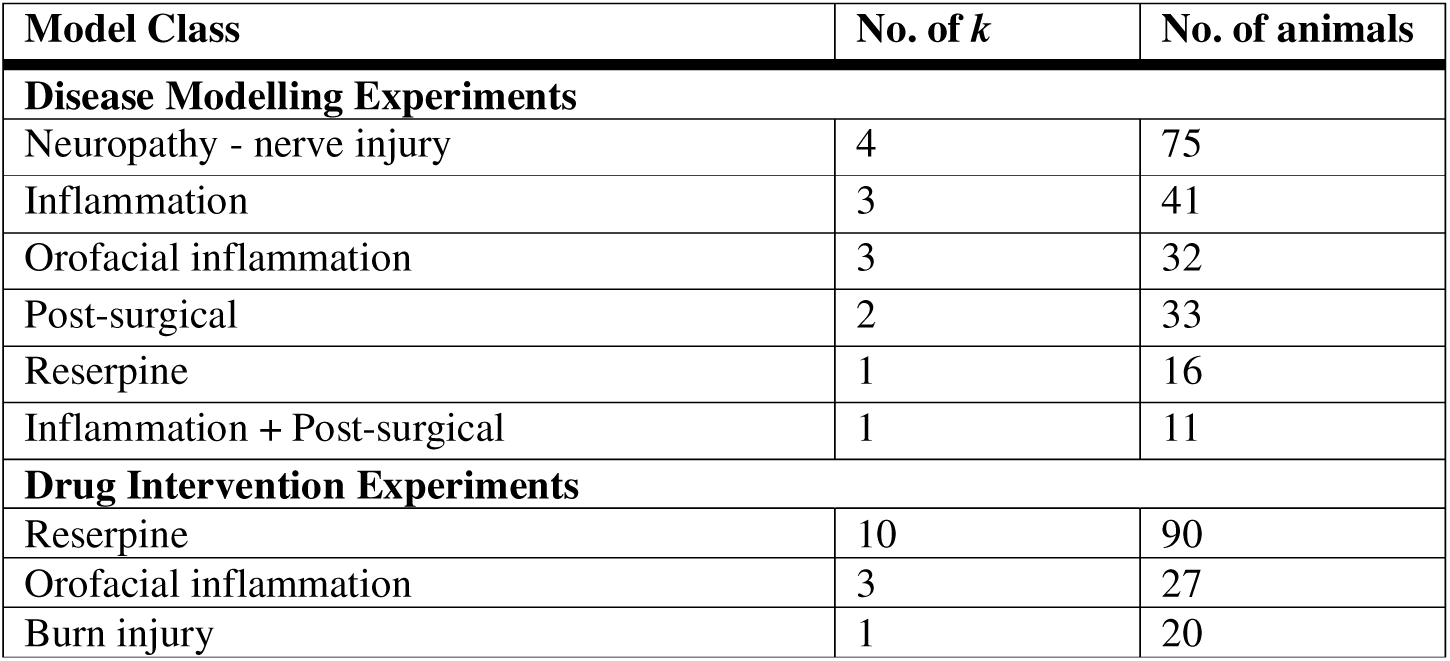
Model classes reported in studies reported both RGS scores and mechanically evoked limb withdrawal outcomes.

| Model Class | No. of *k* | No. of animals |
| --- | --- | --- |
| **Disease Modelling Experiments** |  |  |
| Neuropathy - nerve injury | 4 | 75 |
| Inflammation | 3 | 41 |
| Orofacial inflammation | 3 | 32 |
| Post-surgical | 2 | 33 |
| Reserpine | 1 | 16 |
| Inflammation + Post-surgical | 1 | 11 |
| **Drug Intervention Experiments** |  |  |
| Reserpine | 10 | 90 |
| Orofacial inflammation | 3 | 27 |
| Burn injury | 1 | 20 |

#### Insufficient data to assess relationship between RGS scores and mechanically evoked limb withdrawal outcomes

The analysis showed no significant correlation between RGS scores and mechanically evoked limb withdrawal outcomes in disease modelling experiments (Fig 9A), but there was a significant correlation in drug intervention experiments (Fig 9B). It should be noted that this negative correlation becomes insignificant (*P* = 0.79) when excluding the most severely affected behavioural data point (SMD, RGS = 6.04; mechanical stimulus = −4.93) from the Goder et al. (2021) study [38], where the burn injured rats were treated with sodium channel blockers.

**Fig 9.**
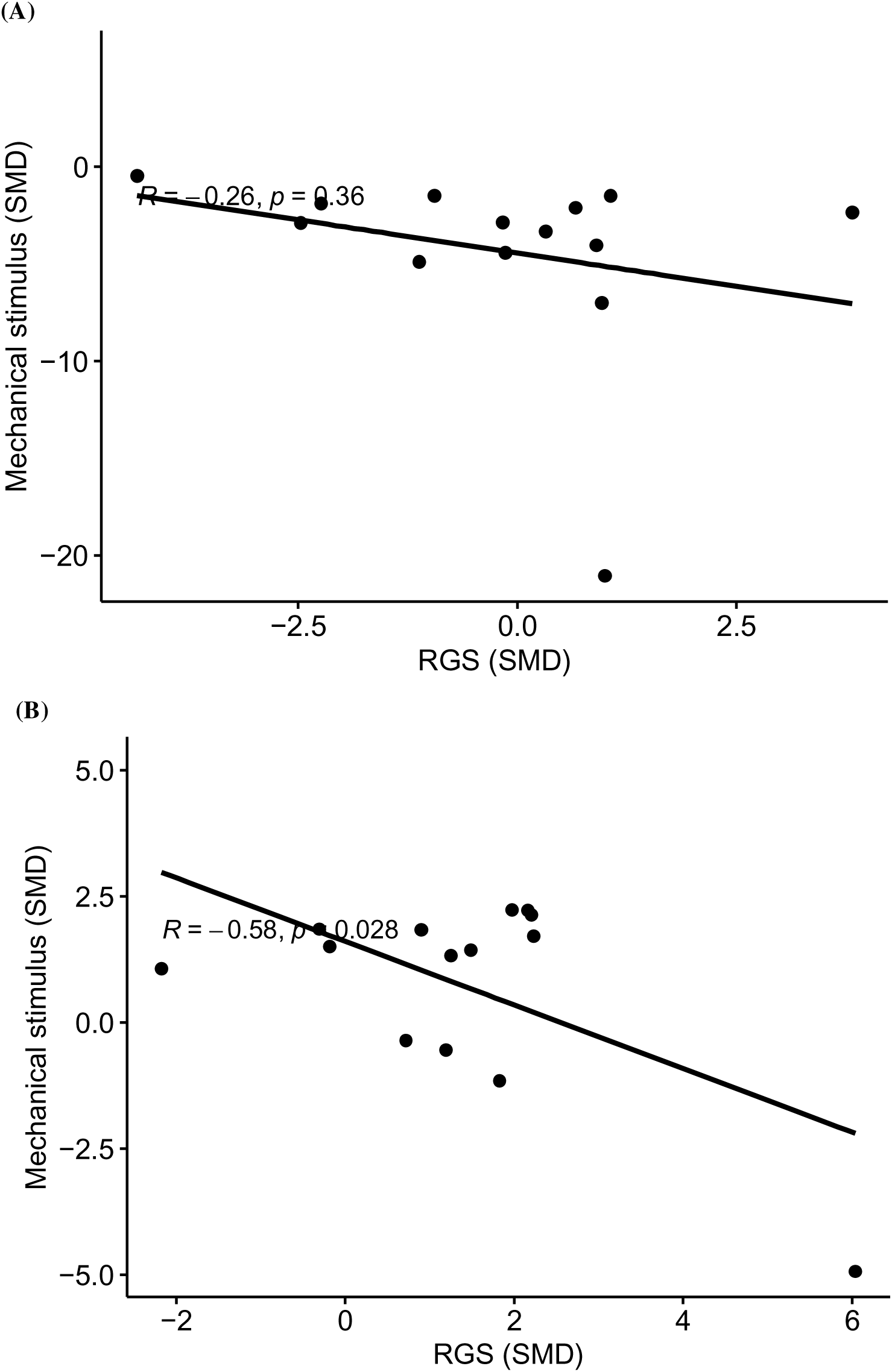
Pearson’s Correlation test of RGS scores and mechanically evoked stimulus outcomes. (A) Rat disease modelling experiments, and (B) drug intervention experiments.

The assessment of the correlation between RGS scores and mechanically evoked limb withdrawal outcomes in rats which received the same treatments was not possible due to lack of data.

#### Risk of bias

The overall risk of bias of the 54 studies is unclear. The reporting of random group allocation and blinding of outcome assessment were moderately high (72%, 39 studies). However, the reporting of other methodological quality criteria was relatively low: predefined animal inclusion criteria (9%, 5 studies), allocation concealment (11%, 6 studies), sample size calculation (20%, 11 studies), and animal exclusion (20%, 11 studies) (Fig 10A). This contrasts with the high reporting of conflict of interest (81%, 44 studies) and compliance with animal welfare regulations (94%, 51 studies). The methods used to mitigate bias were rarely reported hence an unclear risk of bias (Fig 10B). A traffic light plot presenting the risk of bias score for each report is available in S3 Appendix.

**Fig 10.**
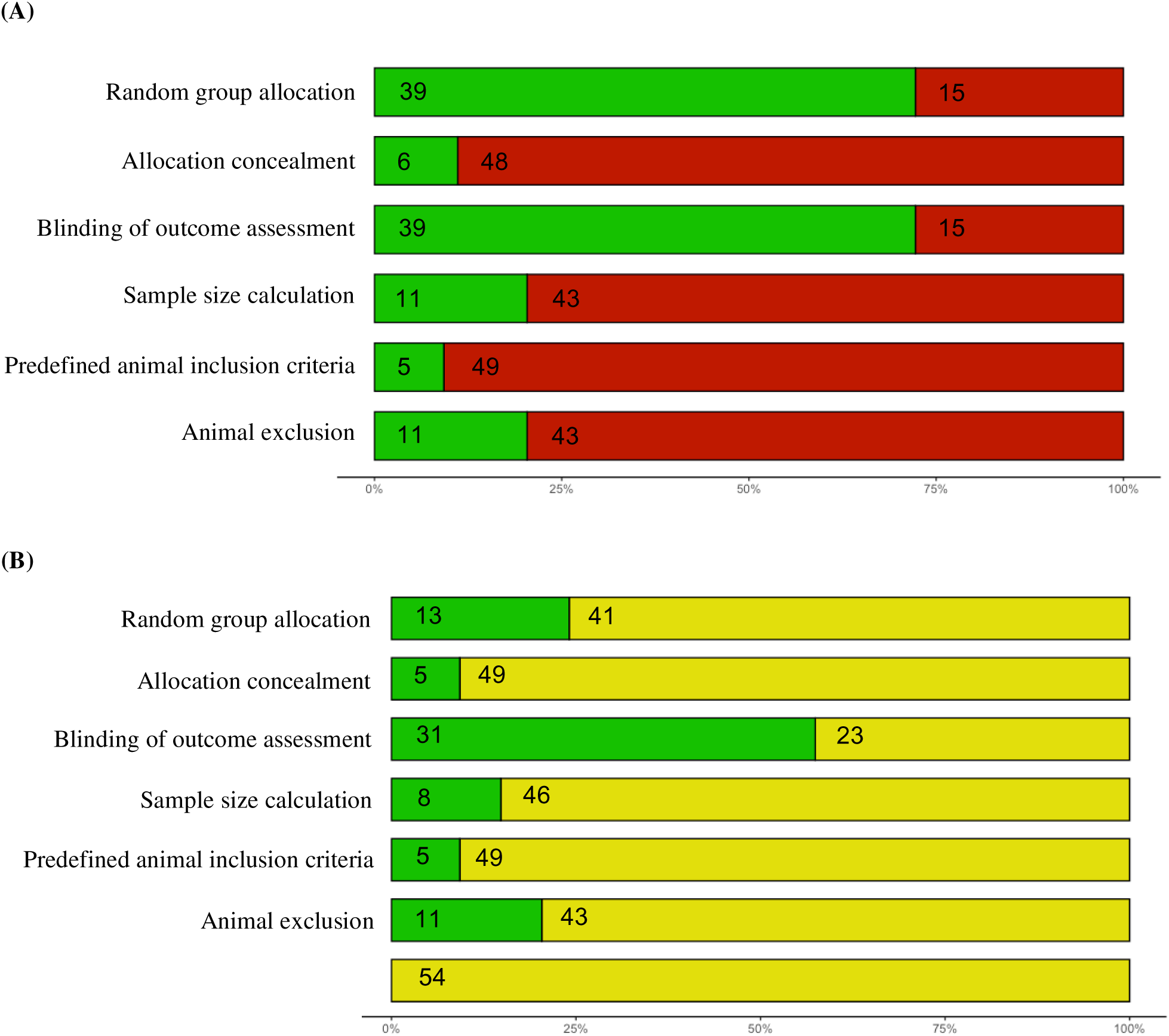
Risk of bias assessment: RGS score studies. Summary plots showing the percentage of the 54 studies that (A) reported the methodological quality criteria and (B) the corresponding risk of bias score given for each methodological quality criterion. Numbers shown within the bar plots indicate the number of studies. Reporting of a statement regarding potential conflict of interests and compliance with animal welfare regulations were extracted, but they were not part of the overall risk of bias.

#### Impact of methodological quality criteria on RGS scores

In disease modelling experiments, RGS score data of all control types were combined. Only the reporting of sample size calculation accounted for a significant proportion of the observed heterogeneity (Fig 11). Smaller effect sizes were observed in experiments that reported sample size calculation (SMD = −1.55 vs −4.33, *Q* = 7.00, *df* = 1, *P* = 0.008).

**Fig 11.**
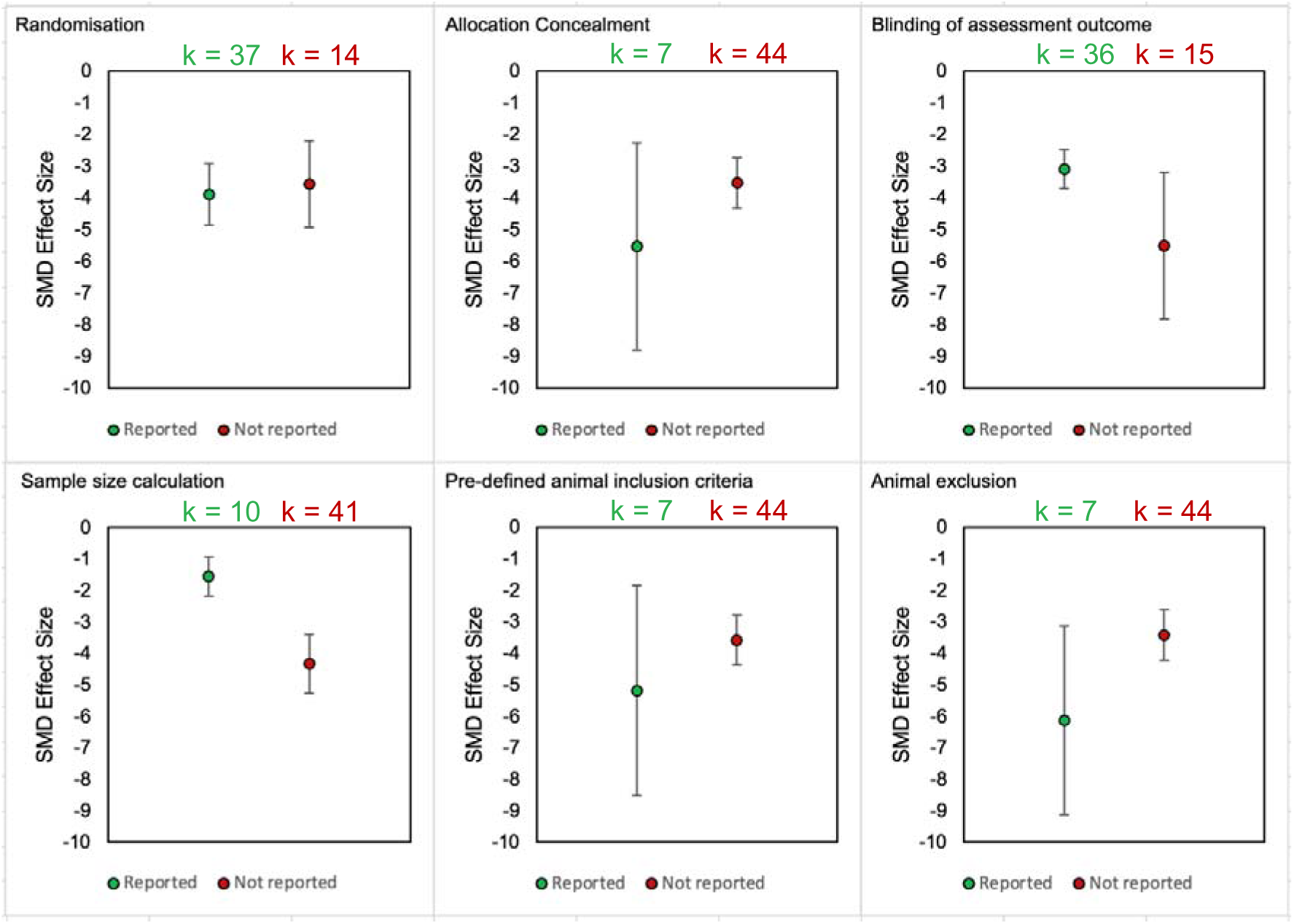
Effect sizes of RGS scores in disease modelling experiments: impact of reporting methodological quality criteria in rats.

In drug intervention experiments, only the reporting of randomisation accounted for a significant proportion of the observed heterogeneity. Larger effect sizes were associated with experiments which reported the reporting of randomisation (SMD = 2.81 vs 0.90, *Q* = 9.00, *df* = 1, *P* = 0.003) (Fig 12). No studies reported allocation concealment therefore we could not assess its influence on RGS effect sizes.

**Fig 12.**
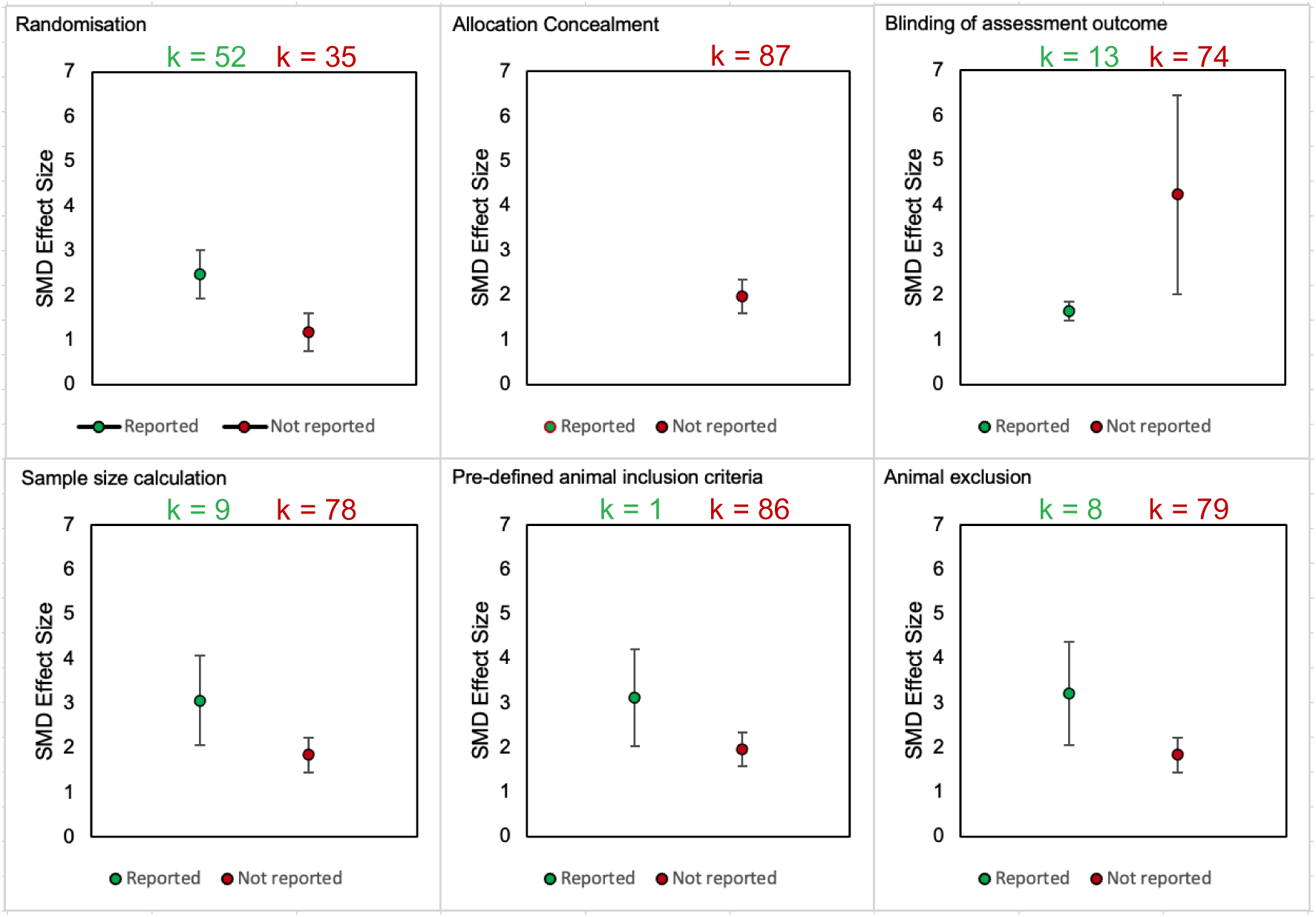
Effect sizes of RGS scores in drug intervention experiments: impact of reporting methodological quality criteria in rats.

#### Reporting quality

All studies were published after the introduction of the ARRIVE guidelines [39]. Out of 54 included studies, 11 studies (20%) stated reporting in accordance with the ARRIVE guidelines, but none of these studies provided a checklist. Furthermore, these studies did not report sufficient details on some of the methods used to mitigate bias. The reporting of methodological quality criteria for these 11 studies was low to moderate (Table 13).

**Table 13.** Reporting of methodological quality criteria of the 11 studies which stated reporting in accordance with the ARRIVE guidelines but did not provide a checklist.

| Methodological quality criteria | No. of studies reported | No. of studies that reported the method | No. of studies that scored low risk of bias | No. of studies that scored unclear risk of bias | No. of studies that scored high risk of bias |
| --- | --- | --- | --- | --- | --- |
| Random group allocation | 9 | 3 | 3 | 8 | 0 |
| Allocation concealment | 3 | 3 | 3 | 8 | 0 |
| Blinding of outcome assessment | 7 | 7 | 7 | 4 | 0 |
| Sample size calculation | 6 | 3 | 3 | 8 | 0 |
| Pre-defined animal inclusion criteria | 2 | 2 | 2 | 9 | 0 |
| Animal exclusion | 2 | 2 | 2 | 9 | 0 |

#### Publication bias

The overall effect size when combining all control types (*i.e., internal, and external controls*) (*k* = 51) is −3.80 [95%CI −5.31 to −2.29]. Egger’s regression test was significant (*P* < 0.001), therefore suggesting the presence of publication bias (Fig 13). Trim-and-fill analysis did not impute theoretically missing experiments.

**Fig 13.**
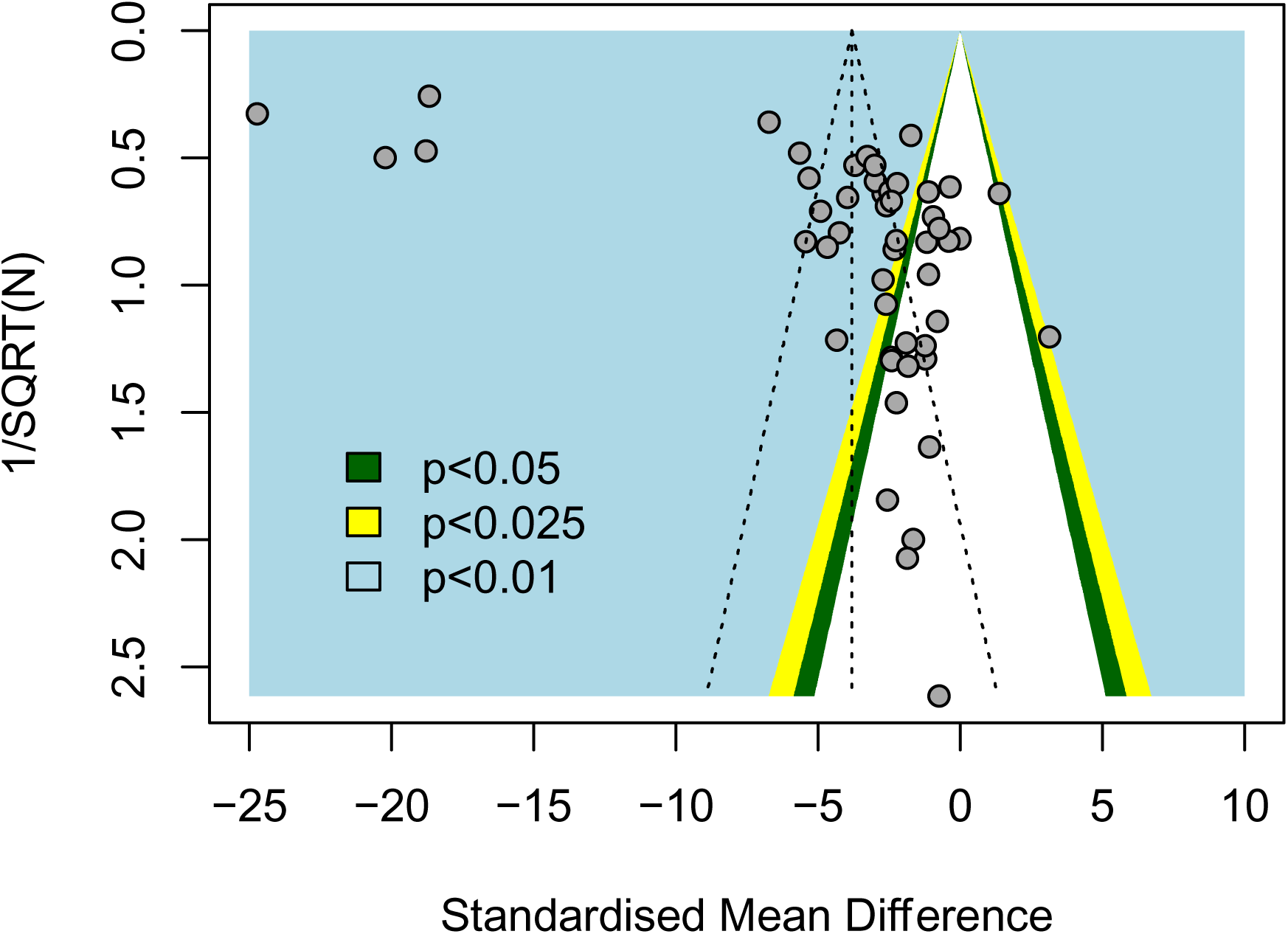
Evaluating publication bias in disease modelling RGS experiments of rats. The vertical dashed line represents the overall effect size. Filled circles represent experiments from the published studies. The coloured backgrounds indicate the statistical significance of effect sizes of cohort-level comparisons.

The overall effect size of RGS scores in drug intervention experiments (*k* = 87) is 1.97 [95%CI 1.25 to 2.69]. Egger’s regression test was significant (*P* < 0.0001), therefore indicating the presence of publication bias (Fig 14). However, trim-and-fill did not impute any theoretically missing data.

**Fig 14.**
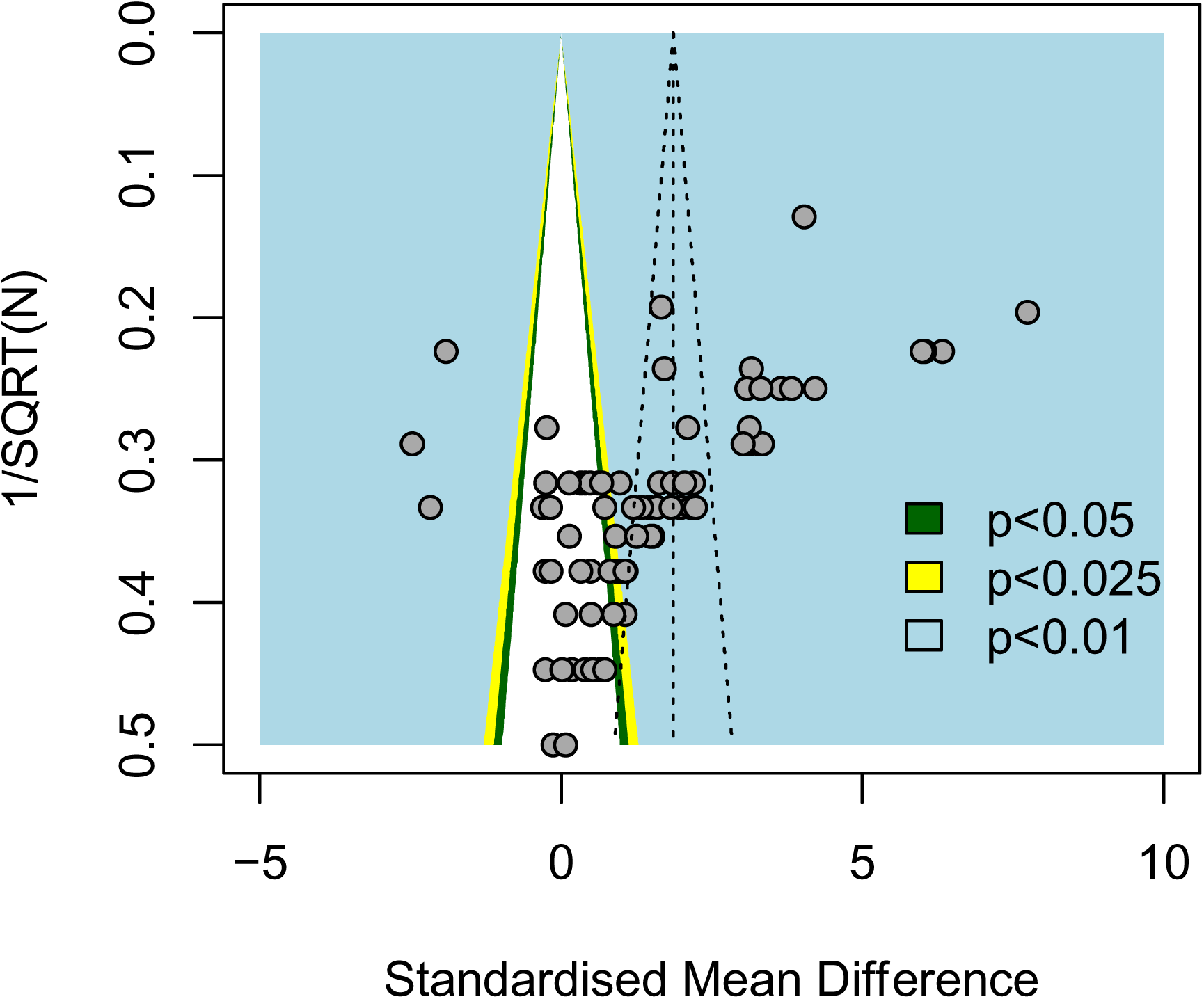
Evaluating publication bias in drug intervention RGS experiments of rats. The vertical dashed line represents the overall effect size. Filled circles represent experiments from the published studies. The coloured backgrounds indicate the statistical significance of effect sizes of cohort-level comparisons.

## Discussion

This systematic review and meta-analysis provide a comprehensive and objective evaluation of published studies exploring how experimental pain affects RGS scores in laboratory rats, as well as the effects of analgesic drug interventions. Additionally, it investigated how different experimental factors influence RGS scores. By summarising the available evidence, this review aims to guide researchers in selecting suitable animal models, outcome measures, and drug interventions for future studies.

In total 54 studies were identified, comprising the effects of 27 persistent pain associated disease models (15 model classes) and 16 drug classes on RGS scores in 1816 rats.

### RGS score is a persistent pain-related behavioural outcome

All included disease models are associated with persistent pain. Their disease onset duration lasted between hours and days, which could be reflected by the median time between model induction and the first and last RGS assessment. The observed increase in overall RGS score in these models, coupled with the attenuation through analgesic drug interventions, suggests that RGS score serves as an acceptable outcome measure for persistent pain. While previous research on mouse grimace scale (MGS) has demonstrated the ability of the rodent grimace scale in identifying pain related to acute nociception lasting from minutes to hours [25], this systematic review could not definitely establish the utility of RGS score as an outcome measure for acute nociception due to insufficient data.

### Effects of animal model characteristics and drug interventions on RGS scores

#### Unable to ascertain the influence of disease models on RGS scores

Twelve classes of rat disease models were reported in disease modelling experiments, but insufficient data for most models prevented stratified subgroup and multilevel analyses. Consequently, the effects of disease models on RGS scores remain unclear, except for orofacial inflammation. While RGS scores were more frequently used to assess pain in rats with orofacial inflammation and post-surgical models, their application in other models, such as those reflecting aspects of neuropathy, spinal cord injury, migraine, and arthritis, was less common. Further studies are required to determine if RGS score is sensitive to changes in less-studied disease models particularly those associated with neuropathy.

### Drug interventions influenced RGS scores

Stratified subgroup analysis suggested that RGS scores were influenced by drugs from the two most frequently reported drug classes: opioids and NSAIDs, primarily in post-surgical models. This aligns with findings from another systematic review that assessed mouse and rat grimace scales in post-surgical models only [40]. A post-hoc multilevel analysis further indicated that drug interventions accounted for 35.14% of the overall heterogeneity, highlighting the impact of drug class on RGS scores.

The reported NSAIDs were diclofenac, ibuprofen, carprofen, ketoprofen and meloxicam. Among these, ketoprofen significantly reduced RGS scores in post-surgical models, supporting its use as a positive control for testing novel drug efficacy in similar models. Current evidence suggests that ketoprofen exhibits superior anti-inflammatory and analgesic effects compared to other NSAIDs such as ibuprofen and diclofenac [41–43], which make it a promising option for managing postoperative pain. NSAIDs are increasingly used as part of multimodal analgesic regimens to minimise opioid consumption [44] and are first-line treatments for inflammatory pain, a major contributor to postoperative pain[45]. We could not ascertain the treatment effects of other NSAIDs due to insufficient data.

The reported opioids included morphine, buprenorphine and fentanyl. Significant improvement of RGS scores were associated with morphine and buprenorphine. However, we could not determine the treatment effects of these three opioids in different model classes due to insufficient data. Fentanyl, studied in a single experiment of in rats modelled with stroke, showed potential but requires further investigation because [46]. current clinical evidence on opioids for post-stroke pain is inconclusive due to either a lack of efficacy or complications associated with drug tolerance at higher doses [47].

Sedation is a common adverse effect associated with opioids, could affect facial expressions scored by RGS. However, its impact on grimace scales remains underexplored. Details such as dose justification, recovery time after surgery, and control for anaesthetic effects, all known to influence RGS [48, 49], were often missing. For example, while RGS was measured 10 hours after model induction in morphine studies, this timeline was not reported in buprenorphine experiments. Hence, controlling these variables is necessary for accurate interpretation of grimace scores.

### Strain did not impact RGS scores

Strain did not significantly influence RGS scores. Sprague Dawley rats, the most commonly reported strain, consistently showed the largest effects of disease modelling and drug treatments. Currently, research on the effects of using different rat strains on RGS scores is limited, with only one study conducting between-strain comparisons in RGS scores [50]. In this study, RGS scores was compared between two female rat strains, Sprague Dawley and Wistar, undergoing laparotomy procedures and subsequent administration of meloxicam and buprenorphine [50]. The study found no significant difference between the strains [50]. This contrasts with observations in MGS scores, where significant differences were identified in the baseline MGS scores of healthy mice across four different strains [30].

### The influence of sex and control types on RGS scores remains uncertain

Due to limited number of studies including female animals, we were unable to determine the effect of sex on RGS scores. Two studies compared RGS score differences between male and female Sprague Dawley rats modelled with meningeal neurogenic inflammation, induced by two different methods, and they showed contradictory findings; one indicated the presence of sex differences in RGS scores [51], while the other did not [52]. However, these studies may not have been powered to assess sex differences [53]. The known influence of sex on drug treatment effects [54–56], as demonstrated in a MGS study [57] underscores the need for more balanced representation of sexes in pain-related RGS research.

The review categorised rat disease modelling experiments based on the type of control used, distinguishing between internal baseline control and external control. However, due to limited data, the review could not compare the effects of using different control types on RGS scores in rats modelled with the same disease conditions, and this is also an area that currently has not been extensively explored. Baseline grimace scores of healthy naïve rats are never at zero [26, 27, 58]. Therefore, it is essential to understand the baseline grimace score in healthy naïve rats and whether this score varies within and between strains, as well as between sexes. However, to date, there have been no studies comparing baseline RGS scores between strains and sexes of laboratory rats. Variability in baseline MGS has been observed among different mouse strains, as well as between sexes within the same mouse strains, although no significant difference was observed in mice from the same strains [30].

### Effects of experimental conditions on RGS scores

The review extracted study characteristics relating to experimental conditions; however, it was not possible to investigate many of their associations with RGS scores because information regarding these characteristics was not reported frequently or in sufficient detail.

### Other considerations relating to rodent grimace assessments Facial action units

Out of the fifty-four included studies, 47 reported using the original four original facial units for scoring, as defined by Sotocinal et al. [26], while the remaining 7 either modified the facial units by excluding specific ones or did not report what units were scored. The most commonly excluded facial unit in RGS was “whisker changes,” a trend similarly observed in a scoping review of MGS [59]. Authors often cited reasons such as inconsistent scoring and difficulties in capturing vibrissae due to poor experimental setups. This represents a feasibility and practical challenge, and we currently do not know the impact of not measuring whisker changes and other facial action unit on the overall score. This challenge could be addressed by the emerging automatic tools developed for objectively scoring facial units in different animal species, using deep learning [60–62].

### Reliability

Out of the 34 studies that reported the number of raters participated in the RGS scoring, 10 studies employed more than one observer, predominantly using two observers. However, within this subset, approximately half did not disclose whether observers rated independently, performed rater agreement analysis, or detailed the reconciliation process for scores. This raises concerns about the reliability of the RGS score data. Currently, the impact of training and experience in using rodent grimace scales on reliability is an under-researched area. Two MGS studies have presented conflicting findings regarding whether training leads to a more reliable assessment of rodent grimace scores. One study suggested that experienced observers exhibited greater consistency [63], while another indicated a good correlation between novice and experienced observers [64].

### Retrospective imaging scoring versus real-time scoring

RGS scoring typically involves retrospective analysis of video recordings, where raters score facial units based on predefined criteria. Real-time scoring, on the other hand, involves continuous observation and scoring of facial units during the event. Only two studies reported real-time RGS scoring in rats, with one focusing solely on whisker changes [65, 66]. The other study compared real-time and retrospective scoring methods [66]. Leung et al. (2016) demonstrated that real-time 15-second interval observations successfully discriminated between control and treatment groups, like the results obtained using the standard retrospective imaging scoring method. Due to limited evidence, this review could not conduct analyses to assess the impact of scoring methods on RGS scores, nor could it evaluate whether real-time scoring can be influenced by the presence of human observers. Such influence has been suggested to cause marked differences between real-time scoring and standard retrospective imaging scoring in MGS, where live scores were significantly lower than the corresponding retrospective imaging scores [30, 67].

### Correlation of RGS scores and mechanically evoked limb withdrawal behaviours

RGS scores were negatively correlated with mechanically evoked limb withdrawal behaviours in drug intervention experiments. As RGS scores improved, the severity of the mechanically evoked outcomes worsens. However, this correlation was primarily driven by a single burn injury experiment, which had the largest treatment effect. Due to limited mechanically evoked data, the relationship between RGS scores and these evoked behaviours remains unclear.

Comparing RGS scores to stimulus-evoked behavioural outcomes may not be meaningful, as they measure different aspects of the pain experience, especially where neuropathic pain is being investigated as this condition more often associated with sensory loss than gain. To enhance the reliability and depth of rodent pain research, we recommend using a combination of stimulus-evoked, non-evoked and ethologically relevant behavioural assessments [8, 9, 68].

### Risk of bias and reporting quality

Randomisation and blinding were the most frequently reported risk mitigation measures, but allocation concealment and pre-defined animal inclusion criteria were the least reported. Drug intervention experiments reporting randomisation were associated with larger treatment effects, while disease modelling experiments reporting sample size calculations were associated with smaller effect sizes. Bias mitigation methods were poorly reported. This hindered accurate assessment of internal validity, leading to unclear risk of bias ratings for most studies. Similar issues were found in other preclinical systematic reviews of pain [7–9]. Few studies adhered to the ARRIVE guidelines [39], and those that did lacked checklists and detailed reporting of methodological quality criteria. Inconsistent methods limit research reliability. Researchers should adopt quality frameworks like EQIPD [69] to improve study design and report in accordance with an established guideline, such as the ARRIVE guidelines 2.0 [32]. In addition to the ARRIVE 2.0 guidelines, we have outlined a list of essential reporting items for RGS assessment in S4 Appendix.

### Publication bias

Our analysis suggests potential publication bias in both animal models and drug treatments, indicated by Egger’s regression test. However, trim and fill did not impute any theoretically missing studies. This may reflect the limitations of the tests; for example, Egger’s regression and trim-and-fill are less reliable for small studies with large between-study heterogeneity [70–73]. Additionally, factors like selective reporting, study quality, between-study heterogeneity can also lead to a funnel plot asymmetry [74].

### Limitations

We conducted comprehensive database searches using search strategies that had a good balance between sensitivity and specificity. While it is possible some relevant studies were missed, searches were all conducted systematically so the included studies represent an unbiased sample.

Our search was conducted on 24^th^ January 2022, but given the substantial size of the review the inclusion of newer studies at this time is unlikely to change the overall conclusions about the use of the RGS in rodent pain research. Nevertheless, future updates will be necessary to incorporate new evidence, which could provide new insights into the sources of heterogeneity and how best to utilise the grimace scale. One option is to transform this review into a “living review” that continuously updates and integrates evidence as it becomes available [75].

The review is limited to the information reported in the included studies. For example, while bias mitigation measures may have been implemented but not reported, some studies might falsely claim to have implemented these measures or might have used inadequate methods. The low prevalence of reported methodological quality measures might have reduced the power of risk of bias assessment, leading to inconsistent relationship between these measures and RGS effect sizes.

Stratified subgroup analysis is limited as it only examines one variable at a time, disregarding other factors such as disease models and drug treatments. Moreover, it cannot pinpoint how much a specific factor contributes to overall heterogeneity. To address this, we conducted *post-hoc* multilevel analyses. However, a limitation of the multilevel analysis is that, after grouping data at few levels, the sample size becomes notably small, compounded by the initial small sample size.

We could not determine how drug administration regimens affect RGS scores due to insufficient data. We also could not compare different animal characteristics within the same disease. Many details about study design, experimental conditions, and assessment characteristics were poorly reported, preventing us from drawing meaningful conclusions. Other variables and factors, like the sedative effects of a drug, were not assessed. Therefore, the results should be interpreted cautiously, with the need for confirmation through prospective experiments.

We extracted behavioural data from the time point showing the greatest difference between treated and untreated animals. This allowed us to assess treatment effects without considering treatment duration but limited our ability to compare different treatment timings. To address this, we also extracted the time between model induction and initial/final rat grimace assessment, as well as the timing of the first drug dose and the start of rat grimace assessment. However, due to significant variations between studies and limited data, we could not analyse the impact of different treatment timings.

Finally, for the meta-analysis, disease models were categorised based on their general pain mechanisms, even though the specific causes might differ. Similarly, drugs were grouped by their primary mechanism of action, disregarding their other properties.

To address the existing gaps and challenges in using the RGS for pain assessment, we propose the following recommendations for future research (Table 14). These recommendations aim to improve the reliability, applicability, and standardisation of RGS in laboratory studies.

**Table 14.** Recommendations for Future Research using the RGS.

| Research Area | Current Gaps | Recommendations |
| --- | --- | --- |
| RGS as a pain-associated outcome measure | Insufficient data on the use of RGS for acute nociception. | Conduct studies to evaluate RGS sensitivity to acute pain |
|  |  | models |
| Disease model influence on RGS scores | Limited data on most disease models except orofacial inflammation. | Expand research on a wider range of models |
| Drug intervention effects | Limited data on NSAIDs and opioids across different models.<br>Limited data on administration regimens. | Further research on the effects of different NSAIDs and opioids across disease models. This should also include a wider range of drugs and impact of side effects e.g., sedation.<br>Study the impact of different drug dosages and administration regimens on RGS. |
| Strain influence on RGS | No significant strain differences were observed, but data is limited. | Conduct more studies comparing RGS across multiple strains. |
| Sex differences in RGS scores | Limited studies with female rats. | Ensure balanced male and female representation in future RGS studies. |
| Baseline RGS variation | No studies comparing baseline RGS between strains and sexes. | Conduct baseline comparisons across strains and sexes. |
| Reliability of RGS scoring | Observer training and agreement impact are underexplored.<br>Limited studies comparing different methods of scoring e.g., real-time vs retrospective and use of automation technologies | Investigate the accuracy and feasibility of RGS scoring. |
| Environmental conditions influence on RGS | Limited reporting of environmental conditions has limited analysis. | Standardise and improve reporting on experimental conditions in studies using RGS. |

## Conclusion

RGS scores are acceptable outcome measures for assessing persistent pain lasting hours to days, but its full utility requires to be confirmed when more data from diverse disease models become available. The review suggested ketoprofen is a potential positive control for assessing RGS scores in post-surgical rat models. To better determine the appropriate use of RGS, studies are required that use female animals, diverse rat strains, and rigorous methods. Detailed reporting will also be essential. Stimulus-evoked tests and RGS scores should be considered as separate tools with distinct purposes, used together for a comprehensive understanding during rodent pain research.

## Associated data and resources

We have made the full data set available on the Open Science Framework (https://osf.io/27awc/). Specifically, the ‘Screening results table’ https://osf.io/ew346 includes all studies identified in the literature search and includes reasons for each excluded study. It also includes the studies considered eligible for the review. Only the eligible studies had data extracted from. The ‘Raw_data_SyRF’ file https://osf.io/56s7k includes the data extracted from each study, the name of the data extractors and the time of data of extraction. Each study was extracted in duplicate with disagreements reconciled by a third independent reviewer. The ‘Reconciled outcome data’ file https://osf.io/tzfd8 contains all data that would be needed to replicate the analysis. Risk of bias assessments are for each study can be found in the following files ‘Risk of bias (Disease Modelling)’ https://osf.io/mwpc6 and ‘Risk of bias (Drug Interventions)’ https://osf.io/mwpc6.

## Acknowledgements

The authors would like to thank Cristina Marin and Kayli Colpitts for their contribution to the data extraction.

## Supporting Information

S1 Appendix - Full annotations used for data extraction

S2 Appendix - RGS score dataset: disease modelling experiments using baseline internal controls

S3 Appendix - RGS systematic review: traffic light plot

S4 Appendix - Essential reporting items for RGS assessment

