## Supplemental Files for "A systematic review and meta-analysis of the effect of experimental pain on rat grimace scale scores"

### S1 Appendix

#### Study Design Tab

1. Is this study relevant for this systematic review? (Yes/No).
   [No] Why should this article be excluded?
   [List] – no multiple selection
   - Not a primary research study
   - Not a rat in vivo study
   - Does not study pain (acute nociception and/or persistent pain-related disease models)
   - Does not conduct the rat grimace score assessment

Further questions for relevant studies (i.e. when selecting "Yes" for Q1)

2. Is there missing information required for meta-analysis? (Yes/No) (allow multiple choices)
   a. [Yes] What information is missing?
      - Aggregated rat grimace score (e.g. no aggregated score reported)
      - Sample size (i.e. N number)
      - Variance (e.g. S.D., S.E.M)

Reporting guideline

[Yes] Was the study reported in accordance with a reporting guideline? (Yes/No)
*Was the study reported in accordance with a reporting guideline?*

a. [Yes] Which reporting guideline was used? (Choose from the list)
   1. ARRIVE
   2. Landis 4
   3. PRECISE
   4. Other => State "other".

b. [Yes] Did the study provide a checklist as evidence of reporting in accordance with the guideline? (Yes/No)
*e.g. Whether the study stated, "a checklist is provided in the supplement".*

Acclimatisation and animal husbandry

4. State the acclimatisation time (h) before experiments start.
   *Time period of acclimatisation to housing environment following transportation. If data is given as a range, the most conservative estimate will be extracted. E.g. x = 3-5, 3 will be extracted.*

5. In which phase, was the RGS assessed?
   - Light
   - Dark
   - NR

6. What was the light intensity in the testing room? (textbox)
   *State the light intensity and its unit. If not described, write "NR".*

7. What was the temperature in the testing room?
8. What was the humidity in the testing room?
9. What was the noise level in the testing room?
10. What was the vibration level in the testing room?

RGS study level questions

11. In which phase, was the RGS assessed?

- Light
- Dark
- NR

12. Was the RGS used as an animal welfare measure? (Yes/No)
i.e., It wasn't used in relation to the experiments but used as an assessment to measure animals' wellbeing.

13. Did the authors classify the experimental model as a type of …?
Name the model in the comment box [multiple selections]
Acute nociception
Persistent pain-related model
NR

[Any selection] Is the RGS assessment sensitive to this model (yes/no)
[any selection] did the authors report this finding? (yes/no)

Risk of bias assessment

14. Do the authors report random allocation to group? (Yes/No)
*Do the authors describe randomising the animals to control and to treatment groups?*

a. [Yes] What randomisation method did the investigator(s) use?
*Copy and paste from the study into the text box the method that the authors describe e.g. random number generator, random number table. State 'not reported' if the method is not described.*

15. Do the authors report allocation concealment? (Yes/No)
*Are the investigators blinded to which group the animals are allocated?*

a. What allocation concealment method did the investigator(s) use?
*Copy and paste or paraphrase from the study into the text box the method that the authors describe. State 'not reported' if not described.*

16. Do the authors report blinded assessment of outcome? (Yes/No)
*Were the investigators conducting the burrowing assessment blinded to which group the animals were (treatment or control)?*

a. What blinding method did the investigator(s) use?
*Copy and paste or paraphrase from the study into the text box the method that the authors describe. State 'not reported' if not described.*

17. Do the authors report a statement of sample size calculation? (Yes/No)

a. What calculation method did the investigator(s) use?
*Copy and paste or paraphrase from the study into the text box the method that the authors describe. State 'not reported' if not described.*

18. Do the authors report to have pre-defined animal inclusion criteria? (Yes/No)

19. Do the authors report animals excluded from the study? (Yes/No)
*Were animals excluded from the study?*

a. [Yes] What were the reason(s) for the exclusion? (Choose from the list – make sure to treat each exclusion as one.. add a + so people can add more than one!)

- Failure to meet pre-defined inclusion criteria
- Surgical complications
- Death
- Other => state other
- NR

i. [Yes] Were the number of animals excluded for this reason reported? (Yes/No)

I. [Yes] How many animals were excluded for this reason?

20. Do the authors provide a statement of a potential conflict of interest? (Yes/No)

21. Do the authors give a statement of compliance with animal welfare regulations? (Yes/No)

**Disease Model Induction Tab**

Control questions:

1. What type of control does this group represent? (Choose from the list)

- External control: sham (prioritised)
- External control: naïve
- External control: wild-type (wt)
- External control: wt littermate
- External control: wt age-matched same strain
- External control: wt age-matched not same strain
- External control: wt same strain (not age-matched)
- Internal control: baseline
- Other (specify in the comment box)

a. [Sham] What type of disease model does this sham control for? (Choose from the list)

  - Acetic acid (writhing test)
  - Antiretroviral
  - Cancer
  - Carrageenan
  - Chemotherapy
  - Chronic constriction injury
  - Complete Freunds Adjuvant
  - Crush
  - Diabetes
  - Formalin
  - Sciatic nerve ligation
  - Spinal cord injury
  - Streptozotocin
  - HIV protein
  - HIV + antiretroviral
  - Traumatic nerve injury
  - Varicella zoster virus
- Collagen induced arthritis

- Medial meniscectomy
- Tibial nerve transection
- Reserpine
- Glyceryl trinitrate
- Lentivirus vector
- Other (specify in the comment box)

b. [Sham] Was the sham procedure identical to the model induction procedure before injury? (Yes/No/NR)

i. [No] Describe the alternative sham procedure.
*Copy and paste or paraphrase from the text.*

c. [Sham] Was perioperative analgesia administered? (Yes/No/NR)

i. [Yes] Was the same perioperative analgesia administered in the sham group? (Yes/No)
*Was the same perioperative analgesia administered in the disease group, given to the sham model control group?*

1. [No] What was the perioperative analgesia administered?
*State the name(s).*

<u>Non-control questions:</u>

1. [Non-control] What is the type of model? (Choose from the list)
    - Acetic acid (writhing test)
    - Antiretroviral
    - Cancer
    - Carrageenan
    - Chemotherapy
    - Chronic constriction injury
    - Complete Freunds Adjuvant
    - Crush
    - Diabetes
    - Formalin
    - Sciatic nerve ligation
    - Spinal cord injury
    - Spared nerve injury
    - Streptozotocin
    - HIV protein
    - HIV + antiretroviral
    - Traumatic nerve injury
    - Varicella zoster virus
    - Collagen induced arthritis
    - Medial meniscectomy
    - Tibial nerve transection
    - Reserpine
    - Glyceryl trinitrate
    - Lentivirus vector
    - Other (specify in the comment box)

22. [Non-control] Was the model surgically induced? (Yes/No)
*Did the animals undergo a surgical procedure to model a disease or cause injury?*

i. [Yes] What method was used to induce the model?

*Copy and paste or paraphrase description from the text e.g. sciatic nerve exposed under anaesthesia and wrapped loosely in cellulose containing 400 ng gp-120 protein.*

v. [Yes] Was anaesthetic used? (Yes/No)

iii. [Yes] Was perioperative analgesia administered? (Yes/No/NR)

I. [Yes] What was the name of the analgesic administered?

II. [Yes] What dose was administered?
*State the dose and its unit e.g. 20 mg/kg*

III. [Yes] How many times was the drug administered?
*State the number of times the drug was administered. State "not reported" if not described.*

3. [Non-control] Was the model pharmacologically induced? (Yes/No)

i. [Yes] What intervention (drug) was used?

ii. [Yes] What dose was administered?
*State the dose and its unit. E.g. 20 mg/kg*

iii. [Yes] What was the route of delivery? (Choose from the list)

- Dorsum of foot
- DRG
- Plantar surface of paw
- Intradermal
- Intramuscular
- Intraperitoneal
- Intrathecal
- Oral
- Subcutaneous
- Tail vein (Intravenous)
- Other (specify in the comment box)
- NR

iv. [Yes] How many times was the dose administered? (textbox)

vi. [Yes] Was anaesthetic used? (Yes/No)

4. [Non-control] Was the model genetically induced? (Yes/No)

ii. What method was used to induce the model?
*Copy and paste or paraphrase description from the text e.g. lentivirus method*

ii. [Yes] [Yes] Was anaesthetic used? (Yes/No)

iv. [Yes] Was perioperative analgesia administered? (Yes/No/NR)

II. [Yes] What was the name of the analgesic administered?

III. [Yes] What dose was administered?
*State the dose and its unit e.g. 20 mg/kg*

IV. [Yes] How many times was the drug administered?

*State the number of times the drug was administered. State "not reported" if not described.*

**Treatment Tab**

Control questions:

1. [Control] What vehicle was administered?

2. [Control] Was the dosing regimen the same as for treated animals? (Yes/No/NR)

*Was the time in relation to model induction, first and last dose, route of delivery and the total number of administrations the same?*

Non-control questions:

1. [Non-control] What was the name of the treatment administered?

*Copy and paste from the study: name of the drug and alternative names if given.*

2. [Non-control] What dose was administered?

*State the dose and its unit, e.g. 20 mg/kg*

3. [Non-control] What was the route of delivery? (Choose from the list)

- Dorsum of foot
- DRG
- Plantar surface of paw
- Intradermal
- Intramuscular
- Intraperitoneal
- Intrathecal
- Oral
- Subcutaneous
- Tail vein (Intravenous)
- Other (specify in the comment box)
- NR

4. [Non-control] What was the total number of administrations? (textbox)

5. [Non-control] Was the first dose administered pre-model induction? (Yes/No)
   a. [Yes] How long before the model was induced was the first dose administered?

   *Give the time in hours (h).*

6. [Non-control] Was the first dose administered post-model induction? (Yes/No)
   a. [Yes] How long after the model was induced was the dose administered?

   *Give the time in hours (h).*

7. [Non-control] When was the last dose administered?

*Give the time in hours (h) since the first dose was administered to the last dose.*

8. Was the drug administered in combination with another drug? (Yes/No)

*If yes, specify the name(s) of the other drug(s) in the comment box and ensure to add another treatment label(s) to give all the relevant information about the other drug(s).*

**Outcome Assessment Tab**

Integrated question by the SyRF platform

1. Outcome measure average type (Mean or Median)

2. Outcome measure error type (S.E.M. or S.D or IQR)

3. Greater is worse [tick]

4. Outcome measure unit (textbox)

What behavioural outcome assessment was used?
- Rat Grimace Scale
- Nociceptive assays

RGS questions

5. Location of the testing environment
- Home cage
- Novel testing cage
- NR

6. Do authors report rater training for the RGS? (Yes/No)

7. Was the RGS rated by […]
   - a single rater
   - multiple raters (two or more)
   - NR

   [Multiple raters] How many raters were involved? (number)
   [Multiple raters] Were RGS scores rated independently? (Yes/No/NR)
   [Multiple raters] Was a rater agreement analysis conducted? (Yes/No/NR)
   [Yes] What was the method? (copy and paste)
   [Yes] Did the analysis indicate raters were in agreement? (yes/no/other) copy&paste the relevant statements in the comment box.

   [Multiple raters] How scores were reconciled?
   - Disagreement was resolved by discussion
   - Average the scores from raters
   - Other
   - NR

8. Method of RGS scoring (list)
- Image RGS scoring (retrospective)
- Real-time RGS scoring
- NR

[Image RGS scoring] Method of frame selection for RGS scoring
- Manual selection (copy & paste the method)
- Automated selection tool (copy & paste the method)
- NR
- Other (specify in the comment box)

[Real-time RGS scoring] Method of real-time RGS scoring (copy and paste)

9. What facial units were scored? [Multiple selections]
- Orbital tightening
- Nose/Cheek flattening
- Ear changes
- Whisker change
- Other (specify in the comment box)

Were all the conventional facial units scored? (Yes/No)
Orbital tightening, nose/cheek flattening, ear changes and whisker changes.

[No] What facial action unit(s) was not scored?
- Orbital tightening
- Nose/Cheek flattening
- Ear changes
- Whisker change

[any selection] What was the reason? [text]

Were facial action units scored between 0 to 2? (Yes/No)
[No] what was the alternative method? [text]

10. How the final score for each facial unit was calculated?
- Average
- Other
- NR

11. How was the final aggregated score calculated?
- Average
- Sum
- Other
- NR

12. What was the habituation time (min)? (textbox)

*State the time in minutes (if data is given as a range, the most conservative estimate will be extracted e.g. x=3-5, 3 will be extracted)*.

13. What was the assessment duration (h)? (textbox)
    *Given the time (h). If not described, put "NR".*

14. What was the time (h) between model induction and the first RGS assessment?
    *Give the time (h) relative to the model induction. E.g. model induced 3 days before the assessment = 72 h.*

15. What was the time (h) between model induction and the last RGS assessment?
    *Give the last time point in hours that the animal was assessed in this behavioural test relative to model induction. E.g. the test was conducted each day for 28 days, it started 5 days after model induction. 28 + 5 = 33 days = 792 h.*

16. How long after the drug treatment started was the first RGS assessment? (textbox)
    *Give the time in hours (h). If not described, put "NR". If no intervention was used, put "NA".*

---------Nociceptive Assay Annotations-----------

- [Nociceptive assays] What nociceptive assessment was used?
    i. von Frey filament
    ii. von Frey electronic
    iii. Randall Selitto test
    iv. Hargreaves test
    v. Hot plate
    vi. Tail flick
    vii. Tail immersion (hot)
    viii. Acetone test
    ix. Cold plate
    x. Tail immersion (cold)
    xi. Tail clip
    xii. Other (specify in the comment box)

[Nociceptive assays] What is the endpoint? (textbox)
e.g. withdrawal threshold etc.

**<u>Cohort Tab</u>**

1) Is this cohort the control group? (Yes/No)
   a. [Yes] Is this a baseline control (Yes/No)
      ***An aggregated baseline grimace score***

   b. [Yes] How many groups of animals does this control cohort serve?
   *How many groups of animals have received different treatments e.g. a single drug is being tested at 3 different doses in 3 separate groups/cohorts of animals. Therefore, the control cohort (vehicle group) serves 3 cohorts.*

a. What is the rat strain? (Choose from the list)
- Sprague Dawley (mainly this)
- Wistar
- Wistar Hannover
- Lewis
- Transgenic
- Other (specify in the comment box)
- NR

I. [Transgenic] What is the gene of interest? (textbox)
Ai) Has the gene been deleted? (Yes/No)
Bi) Has the gene been inserted? (Yes/No)

b. [Yes] What is the name of the animal supplier?

2. What is the sex of this animal cohort? (Choose from the list)
- Male
- Female
- Both

3. Do the authors report the age of the animals before the study started? (Yes/No)
[Yes] Is the age reported as a range or categorically?
I. [Categorical] What is the age (days)?
II. [Range] What is the lower limit (days)?
III. [Range] What is the upper limit (days)?

4. Do the authors report the weight of the animals before the study started? (Yes/No)
    a. [Yes] Is the weight reported as a range or categorically?
        I. [Categorical] What is the weight (g)?
        II. [Range] What is the lower limit (g)?
        III. [Range] What is the upper limit (g)?

### S2 Appendix

<u>RGS score dataset: disease modelling experiments using baseline internal controls</u>

***A.1. Characteristics of disease modelling experiments in RGS studies using baseline internal controls***

A total of 8 studies, containing 15 cohort-level comparisons, 140 rats, and a sample size range from 5 to 35 with a median of 6 animals per group, assessed the effects of 7 classes of disease models associated with persistent pain on RGS scores (Table 1).

**Table 1.** The number of cohort-level comparisons and animals for animal model characteristics used in RGS disease modelling experiments using baseline internal controls.

| | No. of studies | No. of *k* | No. of animals |
|---|---|---|---|
| **Disease Model** | | | |
| Orofacial inflammation | 2 | 3 | 20 |
| Neuropathy – nerve injury | 2 | 2 | 55 |
| Post-surgical | 2 | 2 | 14 |
| Spinal cord injury | 1 | 4 | 24 |
| Meningeal neurogenic inflammation | 1 | 2 | 11 |
| Inflammation | 1 | 1 | 10 |
| Arthropathy | 1 | 1 | 6 |
| **Rat Strain** | | | |
| Sprague Dawley | 4 | 5 | 74 |
| Wistar | 3 | 8 | 54 |
| Holtzman | 1 | 2 | 12 |
| **Sex** | | | |
| Male | 4 | 4 | 68 |
| Female | 4 | 8 | 50 |
| Mixed | 1 | 3 | 22 |

***A.2. RGS scores were increased by disease models associated with persistent pain***

Disease models significantly worsened RGS scores in disease modelling experiments using internal controls (SMD = -1.89 [95%CI -3.33 to -0.45]). Heterogeneity was high ($Q$ = 134.17, $df$ = 14, $P$ < 0.0001, $I^2$ = 89.6%) (Figure 1).

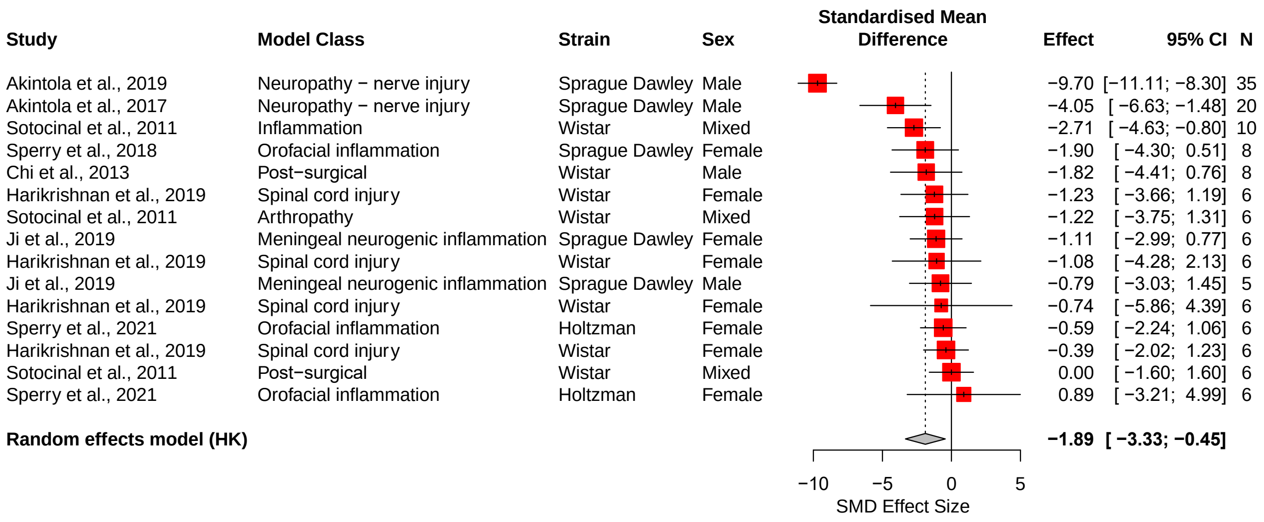

**Figure 1. Effects of disease models on RGS scores: summary forest plot.**
A summary forest plot of the 15 cohort-level comparisons which assessed the impact of disease modelling on RGS scores using baseline internal controls. For each comparison, an effect size was calculated using the Hedges' g SMD method. Effect sizes were pooled using the random effects model.

The restricted maximum-likelihood method was used to estimate heterogeneity. The size of the square represents the weight. CI, confidence interval; N, number of animals.

Stratified subgroup analysis could not be performed due to insufficient data for each characteristic (*i.e.*, $k \geq 10$).

#### S3 Appendix

RGS systematic review: traffic light plot

| Study | Random group allocation | Allocation concealment | Blinding of outcome assessment | Sample size calculation | Predefined animal inclusion criterial | Animal exclusions |
|---|---|---|---|---|---|---|
| Cheng et al., 2019 | Low | Unclear | Low | Unclear | Unclear | Unclear |
| Thomas et al., 2016 | Low | Unclear | Low | Unclear | Unclear | Unclear |
| Lyu et al., 2020 | Low | Unclear | Low | Unclear | Unclear | Unclear |
| Uddin et al., 2019 | Unclear | Low | Low | Unclear | Low | Unclear |
| Tsaousi et al., 2022 | Unclear | Unclear | Low | Unclear | Unclear | Unclear |
| Philips et al., 2017 | Unclear | Unclear | Low | Unclear | Unclear | Unclear |
| Long et al., 2015 | Unclear | Unclear | Unclear | Unclear | Unclear | Unclear |
| Lovrenfcifa et al., 2020 | Unclear | Unclear | Low | Unclear | Unclear | Unclear |
| Sperry et al., 2020 | Unclear | Unclear | Low | Unclear | Unclear | Unclear |
| George et al., 2019 | Low | Unclear | Low | Low | Unclear | Low |
| Gao et al., 2017 | Unclear | Unclear | Low | Unclear | Unclear | Unclear |
| De Rantere et al., 2016 | Low | Unclear | Low | Unclear | Unclear | Low |
| Fujita et al., 2018 | Unclear | Unclear | Low | Unclear | Unclear | Unclear |
| Chagastelles et al., 2020 | Unclear | Unclear | Low | Low | Unclear | Unclear |

| |
|---|
| Low risk of bias |
| High risk of bias |
| Unclear risk of bias |

| | | | | | | |
|---|---|---|---|---|---|---|
| Leung et al., 2019 | Low | Unclear | Low | Unclear | Low | Low |
| Asgar et al., 2015 | Unclear | Unclear | Low | Unclear | Unclear | Unclear |
| Waite et al., 2015 | Unclear | Unclear | Low | Unclear | Unclear | Unclear |
| Kawano et al., 2016 | Unclear | Unclear | Low | Unclear | Unclear | Unclear |
| Akintola et al., 2018 | Unclear | Unclear | Low | Unclear | Unclear | Low |
| Hu et al., 2021 | Unclear | Unclear | Unclear | Unclear | Unclear | Unclear |
| Studlack et al., 2018 | Unclear | Low | Low | Unclear | Unclear | Unclear |
| Goder et al., 2021b | Unclear | Unclear | Unclear | Unclear | Unclear | Unclear |
| Akintola et al., 2019 | Unclear | Low | Low | Unclear | Low | Low |
| Gao et al., 2016 | Unclear | Unclear | Unclear | Unclear | Unclear | Unclear |
| Yamanaka et al., 2017 | Unclear | Unclear | Unclear | Unclear | Unclear | Unclear |
| Harris et al., 2017 | Unclear | Unclear | Low | Unclear | Unclear | Unclear |
| Guo et al., 2019 | Unclear | Unclear | Unclear | Unclear | Unclear | Unclear |
| Sobrinho et al., 2021 | Low | Unclear | Unclear | Low | Low | Unclear |
| Saine et al., 2016 | Unclear | Unclear | Low | Unclear | Unclear | Unclear |
| Thammanichanon et al., 2021 | Unclear | Unclear | Unclear | Low | Unclear | Unclear |
| Akintola et al., 2017 | Unclear | Low | Low | Unclear | Unclear | Unclear |
| Klune et al., 2019 | Low | Unclear | Low | Low | Unclear | Low |
| Goder et al., 2021a | Unclear | Unclear | Low | Unclear | Unclear | Unclear |

| | | | | | | |
|---|---|---|---|---|---|---|
| Sotocinal et al., 2011 | Unclear | Unclear | Low | Unclear | Unclear | Unclear |
| Schneider et al., 2017 | Unclear | Unclear | Unclear | Unclear | Unclear | Unclear |
| Harikrishnan et al., 2019 | Low | Unclear | Unclear | Unclear | Unclear | Unclear |
| Liu et al., 2022 | Unclear | Unclear | Unclear | Unclear | Unclear | Unclear |
| Reed et al., 2020 | Unclear | Unclear | Unclear | Unclear | Unclear | Unclear |
| Whittaker et al., 2016 | Unclear | Unclear | Unclear | Unclear | Unclear | Low |
| Liao et al., 2014 | Unclear | Unclear | Low | Unclear | Unclear | Unclear |
| Chi et al., 2013 | Unclear | Unclear | Unclear | Unclear | Unclear | Unclear |
| Silva et al., 2020 | Low | Unclear | Unclear | Unclear | Unclear | Unclear |
| Guo et al., 2017 | Unclear | Unclear | Unclear | Unclear | Unclear | Unclear |
| Sperry et al., 2018 | Unclear | Unclear | Low | Unclear | Unclear | Unclear |
| Spahn et al., 2017 | Unclear | Unclear | Unclear | Unclear | Unclear | Low |
| Korat et al., 2017 | Unclear | Unclear | Unclear | Low | Unclear | Low |
| Garcia-Robles et al., 2020 | Unclear | Unclear | Unclear | Low | Unclear | Unclear |
| Ji et al., 2019 | Low | Low | Low | Unclear | Unclear | Unclear |
| Korat et al., 2018 | Unclear | Unclear | Unclear | Unclear | Low | Low |
| Koyama et al., 2019 | Unclear | Unclear | Unclear | Unclear | Unclear | Unclear |
| Leung et al., 2016 | Low | Unclear | Low | Low | Unclear | Low |
| Miwa et al., 2019 | Low | Unclear | Low | Unclear | Unclear | Unclear |

| | | | | | | |
|---|---|---|---|---|---|---|
| Iwata et al., 2014 | Unclear | Unclear | Unclear | Unclear | Unclear | Unclear |
| Sperry et al., 2021 | Unclear | Unclear | Unclear | Unclear | Unclear | Unclear |

#### S4 Appendix

**Table. Essential reporting items for RGS assessment.**

| **RGS assessment** |
|---|
| Is it the primary outcome measure? Report the objectives of using RGS scores. |
| Phase of the light/dark circadian |
| Lighting intensity during testing |
| Noise level during testing |
| Habituation time |
| Duration of the assessment |
| Time between model induction/analgesic intervention and the first/last RGS assessment. |
| Presence of human rater. If RGS is scored in real-time, state the number and the sex of the human raters. |
| What are the facial units that will be scored? State the reasons for using an alternative RGS facial units. |
| How many independent raters? |
| Do raters receive training for RGS scoring prior? Report the method (e.g. rater agreement analysis) |
| Is RGS scored using the retrospective image scoring method? Report the method of frame selection, including software version used. |
| Is RGS scored in real-time? Report the method. |
| How is the final aggregated score calculated? |
